# Single-cell profiling of physically interacting immune cells using image-guided sorting

**DOI:** 10.64898/2026.10.09.758094

**Authors:** Ningxin Kang, Cheryl Kim, Thomas J Scriba, Bjoern Peters, Julie G Burel

## Abstract

Single-cell transcriptomics of cell-cell complexes offers key insights into *in vivo* immune interactions, yet distinguishing true physical contacts from artifacts remains challenging. Here, we integrated spectral imaging cell sorting with physical cell deconvolution to separate circulating T cell-monocyte doublets from tuberculosis patients into physically interacting (in-contact) and non-interacting (coincidental) populations. Single-cell sequencing of individual cell partners revealed that physical contact is associated with robust signatures distinct from singlets, including unidirectional monocyte-to-T cell RNA transfer, heightened metabolic activity and T cell cytotoxicity, and a trend toward antigen-specific enrichment. These signatures spanned all T cell and monocyte subpopulations, reflecting a global program of T cell-monocyte engagement. Coincidental doublets displayed intermediate transcriptional profiles between singlet and in-contact states, suggesting the presence of recently disrupted or forming physical pairs. This framework provides high-fidelity deconvolution of cellular interactions, uncovering signatures governing T cell-monocyte communication in human disease.

## Introduction

Intercellular communication governs the coordination and execution of effective host immune responses. Emerging evidence indicates that circulating cell-cell complexes – pairs of immune cells isolated directly from *ex vivo* peripheral blood – represent physically interacting units that reflect ongoing immune dialogue (1). In human peripheral blood, heterotypic T cell-monocyte complexes constitute the most prevalent and widely documented doublet configuration. The frequencies of circulating T cell-monocyte complexes fluctuate dynamically across diverse clinical settings, including acute viral and bacterial infections such as tuberculosis (TB) (2), dengue fever (2), and COVID-19 (3,4), as well as post-vaccination models including influenza (5) and Tdap (2). Furthermore, an increased prevalence of these complexes has been noted in chronic inflammatory states, including HIV infection (6) and obstructive sleep apnea (7).

Beyond simple fluctuations in abundance, circulating T cell-monocyte doublets exhibit molecular characteristics of biological interactions. Microscopic evaluation has revealed features of the classical immunological synapse, including the localized polarization of adhesion molecules (such as LFA-1 and ICAM-1) (2) and accumulation of CD3 at the cell-cell interface during active infection (8). Transcriptomic profiling of these complexes has consistently identified the upregulation of genes associated with T cell receptor (TCR) signaling, major histocompatibility complex class II (MHC-II) antigen presentation, and heightened activation states relative to *in silico* generated doublets or matching circulating singlets (4,6,8). Additionally, these interactive signatures are highly enriched within cytotoxic T cell lineages (4,8). Together, these observations underscore the biological significance of circulating cell-cell complexes as real-time readouts of systemic immune activation.

Despite their clear biological relevance, the study of ex vivo immune cell doublets is severely constrained by persistent technical bottlenecks in their isolation. Heterotypic doublets are typically defined by the dual expression of lineage-specific surface markers via conventional flow cytometry or co-expression of distinct cell-type gene modules in single-cell RNA sequencing (scRNA-seq) datasets. However, these strategies lack the resolution required to differentiate genuine, physically interacting cell complexes from doublet artifacts. These artifacts arise when two independent cells coincidentally pass through the cytometer interrogation point simultaneously, or when two cells are randomly co-encapsulated within the same microfluidic droplet in droplet-based sequencing. Compounding this issue, circulating single cells that abnormally express dual-lineage markers on their surface can appear phenotypically identical to true doublets (9). These technical limitations introduce substantial background noise into downstream analyses; for instance, our recent transcriptomic study of thousands of circulating T cell-monocyte complexes isolated based on CD3 and CD14 co-expression revealed that robust interaction-driven differences resided predominantly within two atypical dual-lineage clusters that comprised less than 20% of the total sorted events (8). The recent emergence of real-time, image-enabled cell sorting represents a paradigm shift for doublet isolation, enabling high-throughput isolation based on instantaneous extraction of morphological and spatial fluorescence features at thousands of events per second (1,10,11). This technology offers a robust solution for eliminating coincidental artifacts prior to downstream analyses such as single-cell sequencing.

In addition to isolation challenges, dissecting the dual transcriptomic signatures of cell-cell complexes introduces significant analytical bottlenecks. Researchers must deploy either bioinformatic or physical deconvolution frameworks (1). Bioinformatic deconvolution algorithms, such as PIC-seq (12), utilize computational modeling of scRNA-seq profiles to infer cell-cell interactions by mathematically decomposing mixed RNA signals back into individual cellular components. While powerful, this approach is fundamentally restricted to broad cell type identification and cannot definitively assign shared metabolic or functional processes to a specific interacting partner. To circumvent this, we recently developed a high-throughput physical deconvolution workflow that couples fluorescence-activated cell sorting (FACS) isolation with droplet-based sequencing (8). Although this method effectively dissociates the cellular partners during sorting – thereby sacrificing direct cellular partner pairing – it yields a high-resolution transcriptomic readout of the individual cells that were recently engaged in physical contact.

To overcome the combined limitations of doublet isolation and transcriptomic deconvolution, we integrated the BD FACSDiscover™ S8 spectral imaging cell sorter (10) into our physical cell deconvolution workflow (8). Our recent work demonstrated the unique power of spectral imaging cytometry to resolve immune cell doublet populations and uncover disease-specific alterations that remain hidden by conventional flow cytometry alone (13). Specifically, we identified a distinct, tight synaptic T cell-monocyte doublet phenotype that is significantly enriched in the peripheral blood of patients with active tuberculosis (ATB) at diagnosis relative to healthy controls (13). Here we sought to further characterize the transcriptomic profile of this synaptic population.

By combining real-time morphometric profiling with spatial fluorescence correlation, we successfully segregated circulating T cell-monocyte complexes into highly refined “in-contact” (physically interacting) or “coincidental” (proximal but non-interacting) categories. Applying this workflow to a cohort of patients with ATB, we isolated thousands of doublets by image-guided sorting for single-cell transcriptomic profiling. Our findings revealed that in-contact doublets harbor distinct, *bona fide* interaction signatures characterized by heightened oxidative phosphorylation, T cell cytotoxicity, antigen-specific enrichment, and unidirectional RNA transfer from monocytes to interacting T cells. Furthermore, high-dimensional clustering demonstrated that these in-contact programs are universally shared within T cell and monocyte subpopulations. Finally, transcriptomic analysis of coincidental doublets revealed a mixed state, demonstrating that some coincidental events retain residual interaction signatures from physical pairs disrupted prior to detection. Together, this work establishes a high-fidelity, image-guided framework to decode genuine intercellular contact in biological samples.

## Results

### Real-time imaging parameters resolve heterogeneity within circulating T cell-monocyte doublets

To dissect the phenotypic morphology of circulating T cell-monocyte complexes, we integrated high-parameter real-time imaging sorting into our established physical deconvolution workflow (8) (**Figure 1A**). This pipeline comprises four sequential steps: 1) identification of CD3^+^CD14^+^ T cell monocyte doublets using multiparametric imaging and fluorescence parameters on a spectral imaging cell sorter; 2) bulk sorting of target populations; 3) systematic disruption of interacting cell-cell complexes into single-cell suspensions via physical shear forces applied during the cell sorting process; 4) droplet-based scRNA-seq combined with TCR sequencing to profile individual interacting partners.

**Figure 1:**
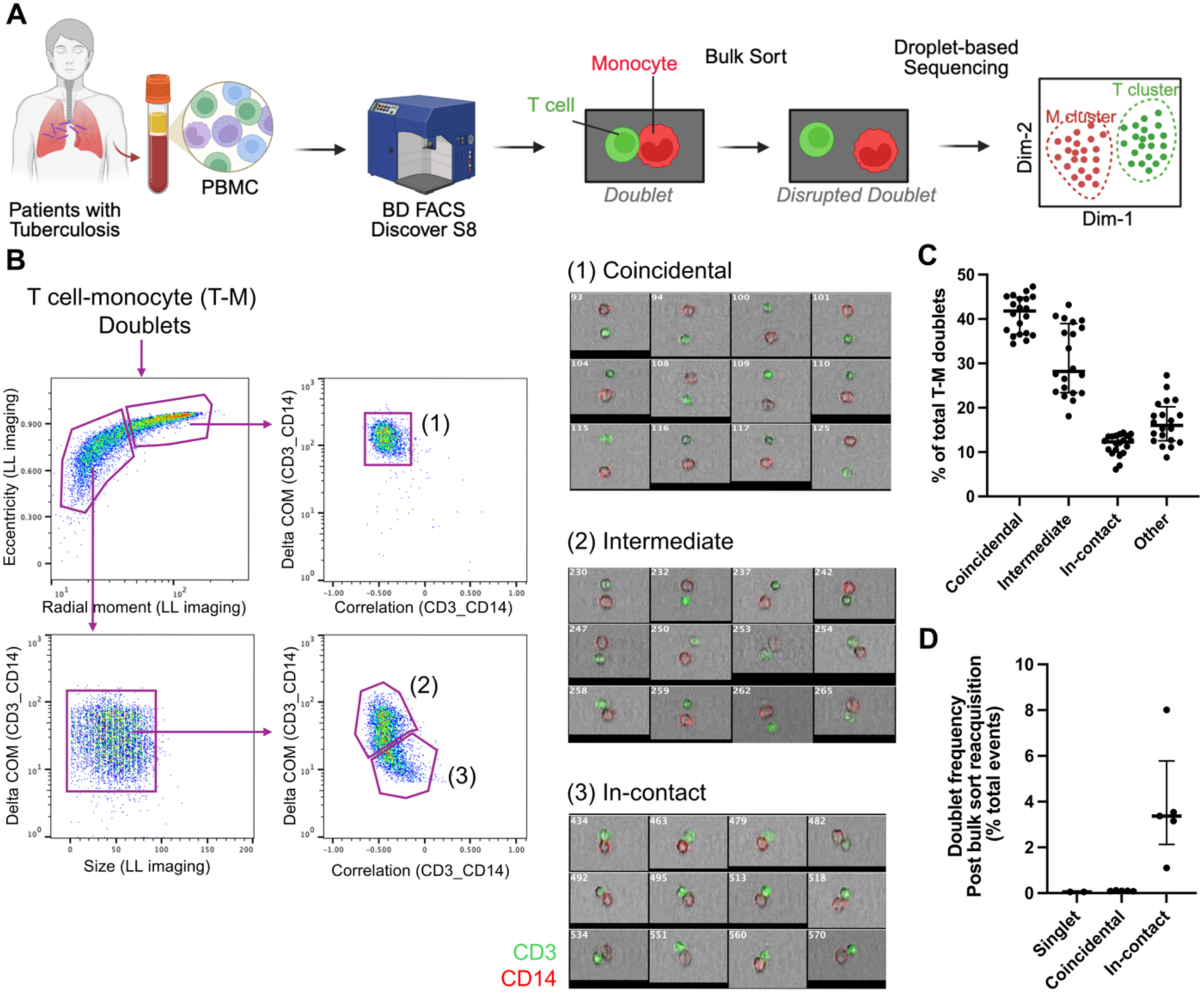
Identification and isolation of distinct T cell-monocyte doublet imaging phenotypes in tuberculosis using imaging cell sorting. (A) Schematic workflow for the isolation and single-cell transcriptomic analysis of T cell-monocyte doublets using the imaging cell sorting technology (BD FACS Discover™ S8). (B) Gating strategy and representative image galleries of different T cell-monocyte doublet phenotypes. T cell-monocyte doublets were initially gated as indicated in Figure S1A. (C) Frequency of different doublet phenotypes in peripheral blood mononuclear cells (PBMC) of patients with ATB collected at diagnosis (n=20). The “Other” category comprises the remaining events not included in the first three categories, representing multicellular aggregates or doublets composed of a CD3^+^CD14^+^ cell and a non-T/non-monocyte (CD3^-^CD14^-^) partner. (D) Frequency of events in a doublet configuration after sorting, determined by re-acquisition on the imaging cell sorter (n=5 PBMC samples).

Using a four-color panel, CD3 and CD14 were captured simultaneously by imaging and spectral detectors, along with CD19 and a viability dye restricted to spectral channels. Doublet events were initially isolated from live leukocytes using eccentricity and radial moment metrics derived from the Side Scatter (SSC) imaging channel – which functions as a darkfield microscopy equivalent (**Figure S1A**). Within this morphometrically defined doublet gate, heterotypic T cell-monocyte complexes were isolated based on co-expression of CD3 and CD14 and absence of CD19 (**Figure S1A**). Visual inspection of image galleries within the CD3^+^CD14^+^CD19^-^ doublet gate revealed marked heterogeneity, ranging from coincidental doublets (non-contacting cell pairs within the same optical field) to true physical doublets displaying either deep synaptic interfaces (e.g., events #69 and #78) or minimal contact areas (e.g., events #75 and #88, **Figure S1B**).

To systematically resolve this heterogeneity, we developed a multi-parametric imaging gating strategy leveraging LightLoss (a brightfield microscopy equivalent), CD3, and CD14 parameters (**Figure 1B**). Coincidental doublets (Population 1) were defined by elevated LightLoss eccentricity/radial moment, low CD3/CD14 fluorescence spatial correlation, and large CD3/CD14 delta center of mass (COM) distances. Within the remaining events, multicellular aggregates and dual-expressing single cells were excluded using LightLoss size and CD3/CD14 delta COM parameters, respectively. Finally, CD3/CD14 correlation and delta COM parameters separated “in-contact” doublets (Population 3) from an intermediate population of adjacent or minimally contacting pairs (Population 2; **Figure 1B**). Thus, real-time imaging parameters resolve extensive morphological heterogeneity within circulating leukocyte complexes, providing a robust framework to isolate highly pure, physically interacting heterotypic doublets.

### Identification and isolation of distinct T cell-monocyte doublet imaging phenotypes in tuberculosis

To characterize the transcriptomic profiles of these imaging phenotypes, we applied our gating strategy to peripheral blood mononuclear cells (PBMC) from a cohort of patients with ATB (n=20). Samples were collected during acute infection immediately following diagnosis and prior to treatment initiation – a sampling window corresponding to peak transcriptomic divergence between doublets and singlets (8). Samples were processed across two independent experiment runs (n=10 each).

Frequencies of T cell-monocyte doublet categories remained remarkably consistent across participants (**Figure 1C**). Coincidental doublets were the most prevalent fraction (median: 41.8%), followed by intermediate doublets (median: 28.2%). In-contact doublets represented the rarest and least variable population, maintaining a tight median frequency of 12.3% across the cohort. The remaining events (”Other” category) comprised multicellular aggregates or doublets composed of a CD3^+^CD14^+^ cell and a non-T/non-monocyte (CD3^-^CD14^-^) partner. Up to 3,000 events each of in-contact doublets, coincidental doublets, and an equal mix of control singlets (T cells and monocytes) were sorted per participant.

Post-sort re-acquisition confirmed that fluidic shear forces efficiently dissociated cell-cell complexes, leaving only a minor residual intact fraction in the in-contact sorted population (median: 3.4%, range: 1.1%–8.0%; **Figure 1D**). Doublet frequencies in post-sort coincidental and singlet suspensions were negligible (medians: 0.09% and 0.05%, respectively; **Figure 1D**). Dissociated single-cell suspensions from each imaging phenotype were immediately processed for droplet-based scRNA-seq.

### Computational identification and removal of inter- and intra-sample doublets

We next implemented a stringent computational pipeline to purge doublets from the scRNA-seq dataset. This rigorous filtering step was necessary due to two confounding factors: 1) droplet-based microfluidics inherently generate stochastic dual-cell encapsulations; and 2) a minor fraction of in-contact doublets resist sorting-induced dissociation and enter the microfluidic chip intact (**Figure 1D**). Consequently, singlet and coincidental doublet libraries are expected to contain two types of doublets: inter-sample and intra-sample artificial doublets (**Figure S2A**, *left*). Sorted in-contact libraries contain a third distinct doublet category: true intra-sample biological doublets that survived fluidic sorting intact (**Figure S2A**, *right*). Although this latter category contains valuable biological information, they will confound our downstream analysis pipeline which assumes pure single cell populations and were set aside for separate, dedicated analyses. Reflecting the presence of these unfiltered doublets, UMAP visualization of the raw data showed a high frequency of dual-expressing cells forming a transcriptomic bridge that blurred the boundary between T cell and monocyte clusters across all library types (**Figures S2B, S2C**).

Our multilayered doublet elimination strategy is schematized in **Figure 2A**. Inter-sample doublets – formed when cells from different participants are co-encapsulated into one oil droplet – were directly identified by tracking multiplexed sample hashtags using a standard hashtag demultiplexing algorithm. Multiplexed droplets containing two hashtags were classified as inter-sample doublets, representing 5.8% of total events (**Figure 2B**). Droplets with zero or more than two hashtags were labeled as empty droplets or higher-order aggregates, respectively, and excluded to retain only singlets (*i.e*., 1 tag assigned, **Figure 2B**).

**Figure 2:**
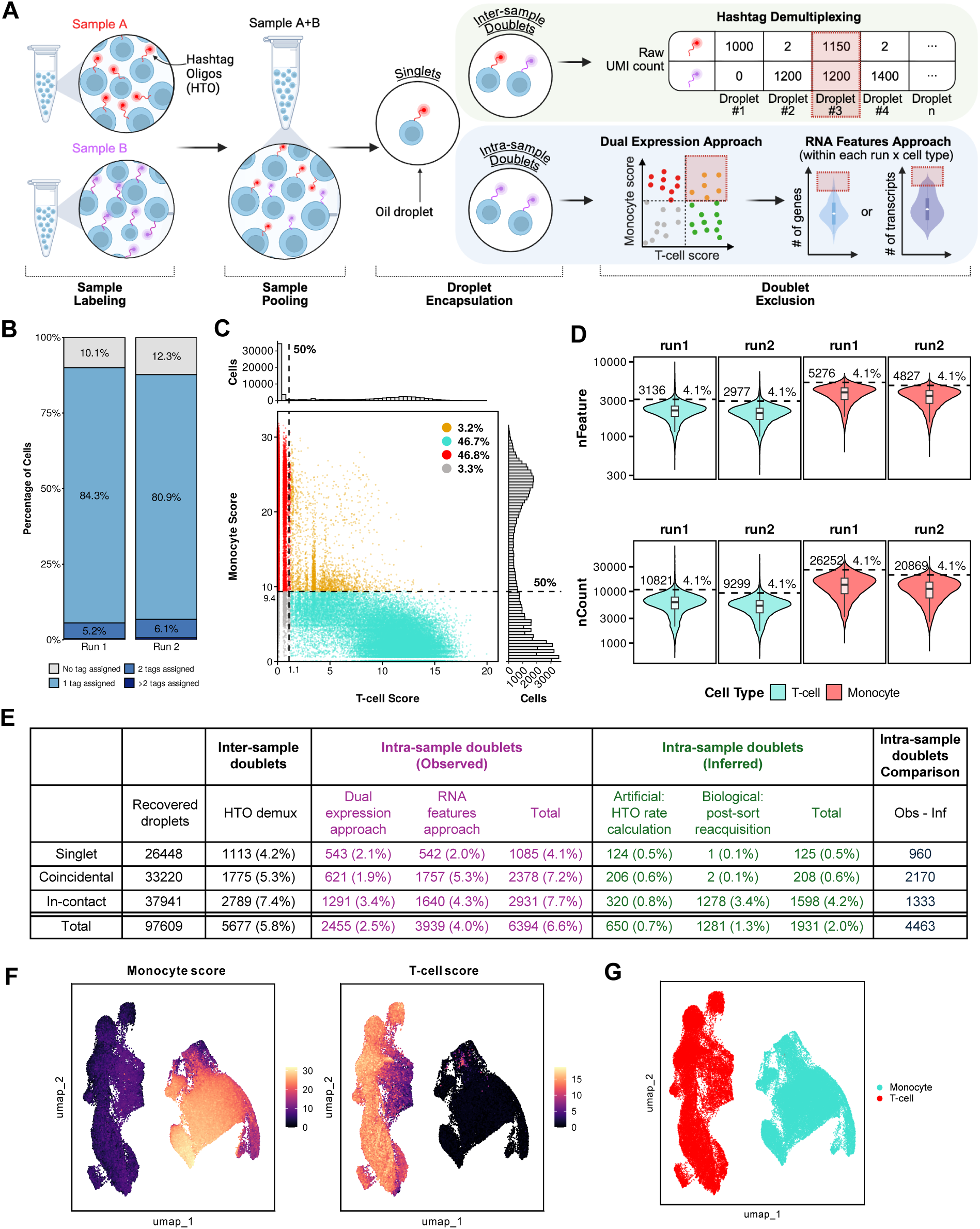
Computational identification and removal of inter- and intra-sample doublets. (A) Schematic depicting the generation of inter- and intra-sample doublets during single-cell droplet-based sequencing, and the three approaches to remove them: hashtag demultiplexing, dual expression approach, and RNA features approach. (B) Stacked bar plot displaying the proportion of cells categorized by HTO demultiplexing sample tag assignment across two independent experimental runs (normalized to 100%). Numerical labels indicate the percentage of total droplets within each category. Data derived from 30,901 droplets from run 1, and 66,708 droplets from run 2. (C) Scatter plot of T cell (x-axis) and monocyte (y-axis) module scores. Dashed black lines indicate 50^th^ percentile thresholds for each score (1.1 for T cells; 9.4 for monocytes). Side histograms display the frequency distributions of T cell (top) and monocyte (right) module scores across all droplets. (D) Violin plots showing the number of genes (nFeature, upper) and transcripts (nCount, lower) distribution per droplet across two sequencing runs (run 1, run 2) and two major cell lineages (T cell, monocyte). Black dashed lines represent the upper outlier cutoff used to remove suspected doublets. Numbers annotated along the dashed line indicate the exact numeric cutoff values, while percentage labels above the lines represent the proportion of outlier droplets removed within each respective group. Inner white boxplots indicate the median and interquartile range (IQR). (E) Summary table of recovered droplet counts alongside observed (HTO-demultiplexed, dual expression, and RNA features) and inferred (artificial and biological) doublets across gating categories. Obs - Inf indicates the difference between observed and total inferred intra-sample doublets. Post-QC uniform manifold approximation and projection (UMAP) plots displaying (F) T-cell and monocyte module score intensities and (G) cell lineage annotation (T cells, turquoise; monocytes, red). For B, C, D, and E, data derived from 76,918 pre-QC droplets; For F and G, data derived from 69,640 post-QC droplets; for all, data derived from n=20 PBMC samples of patients with ATB collected at diagnosis.

Because intra-sample doublets originate from the same participant and cannot be demultiplexed by hashtags, we deployed two sequential approaches to identify and remove them: dual-expression scoring for heterotypic doublets, followed by RNA feature outlier detection for homotypic doublets (**Figure 2A**). For the dual-expression approach, we calculated independent, T cell and monocyte module scores for every droplet; droplets displaying co-expression of both lineage signatures were flagged and removed (**Figure 2C**). Next, homotypic doublets were identified and purged by detecting extreme upper-tail outliers in total unique molecular identifier (UMI) (nCount_RNA) and gene counts (nFeature_RNA). Due to baseline differences in RNA content between lineages and sequencing runs, the upper threshold was defined by a global top quantile rather than an absolute cutoff, with the corresponding thresholds computed separately for each lineage (i.e., T cell and monocyte) and sequencing run. To choose this quantile, we scanned values from 0% to 5% and, for each candidate, compared the number of cells removed to the number of intra-sample doublets estimated (**Figure S2D**). The global quantile was selected as 4.1% as it minimized this discrepancy, and droplets exceeding the threshold in either RNA feature metric were removed (**Figures 2D**, **S2D**, **S2E**). As expected, the dual-expression pipeline specifically eliminated droplets spanning the inter-lineage transcriptomic space, whereas the RNA feature filters excised hyper-transcriptive events within core lineage clusters (**Figure S2F**). Total doublet removals per filtering stage are summarized in **Figure 2E**.

### The computational doublet purging pipeline generates clean cell lineage isolation

To validate the accuracy and stringency of our filtering pipeline, we compared the expected number of inter- and intra-sample doublets against the actual number of events removed from each library type. We observed a linear relationship between observed and expected inter-sample doublet frequencies, which yielded a library-specific doublet factor (DF) (**Figure S2G**). Assuming a constant co-encapsulation rate for inter- and intra-sample doublets, the DF was used to infer the expected number of intra-sample artificial doublets per library (**Figure S2G**). The expected number of intra-sample biological doublets was directly derived from our post-sort re-acquisition metrics (3.4% for in-contact libraries, and <0.1% for coincidental and singlet libraries; **Figure 1D**). The total predicted intra-sample doublet count was consistently lower than the actual number of events excised by our combined dual-expression and RNA feature filtering steps (**Figure 2E**, column 10), confirming that our filtering thresholds were sufficiently stringent to eliminate all residual doublets.

Following doublet removal, low-quality events were cleared using standard lower thresholds gene counts and UMI filters (**Figures S2H**, **S2I**). UMAP visualization of the finalized, clean dataset yielded two cleanly segregated cell islands with no residual transcriptomic bridging in both singlet and doublet categories (**Figure 2F**, **S2J**). These islands were unambiguously assigned as T cells or monocytes based on canonical marker expression (**Figure 2G**) and split into independent data objects for high-resolution downstream analysis of each interacting cell partner.

### The presence of monocyte-specific transcripts in in-contact T cells suggests material transfer

Differential gene expression (DE) analysis between in-contact and singlet T cells identified 90 upregulated (enriched for antibacterial response, cytotoxicity, and oxidative phosphorylation) and 47 downregulated genes (enriched for transcription and adhesion; **Figures 3A**, **3B**, **Table S1**). Surprisingly, several top upregulated genes were canonical monocyte markers not typically expressed in T cells (**Figure 3C**). While the expression of these monocyte-specific genes was elevated in in-contact T cells relative to singlets, it remained significantly lower than the levels observed in the dual-positive (DP) T cell-monocyte droplets previously excluded with our computational filtering pipeline (**Figure 3D**), indicating these events are single cells and not T cell-monocyte doublets.

**Figure 3:**
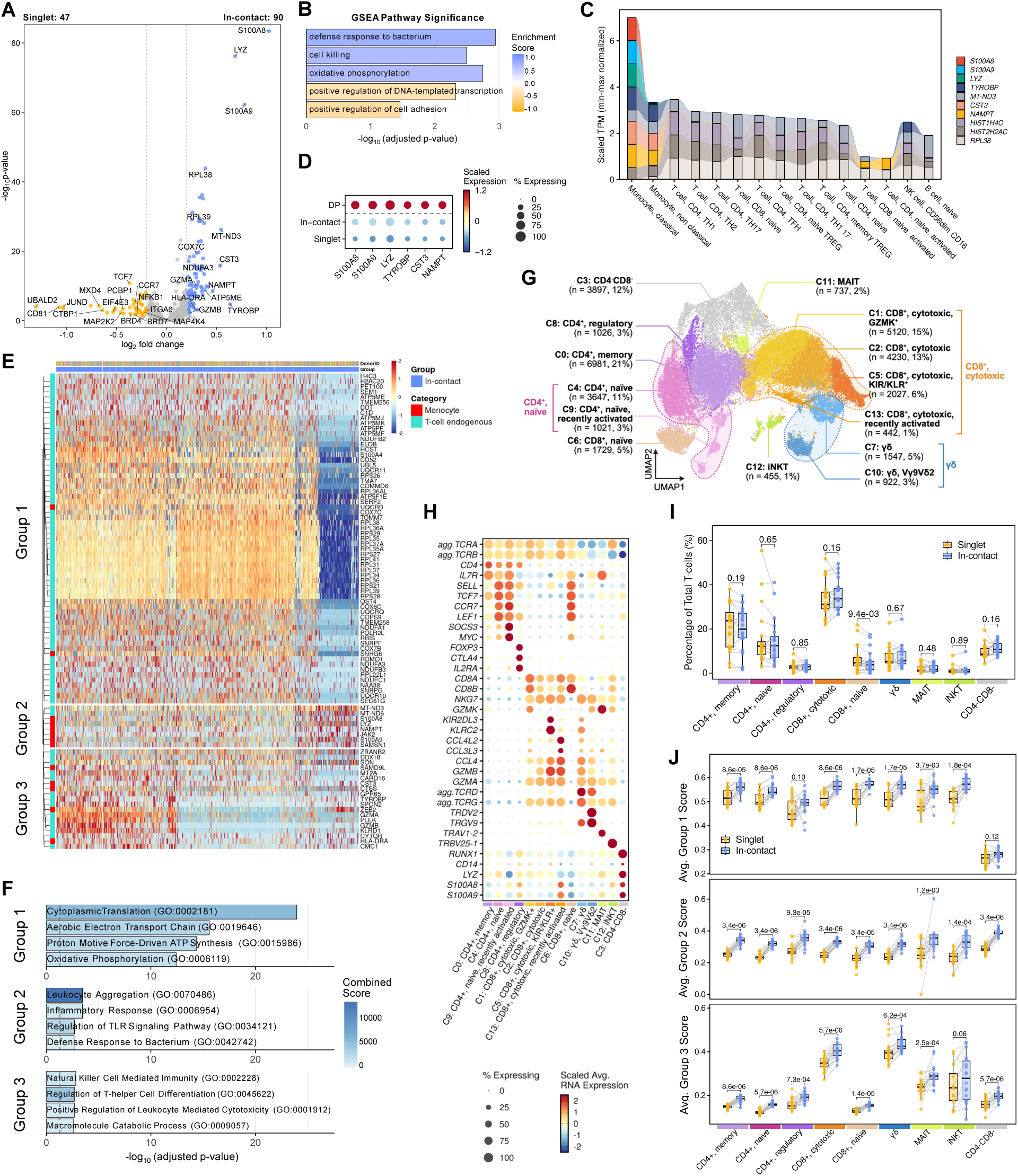
In-contact T cells are associated with specific gene expression signatures. (A) Volcano plot of differentially expressed genes (DEGs) comparing in-contact versus singlet T cells. Genes with Benjamini-Hochberg (BH) adjusted p-value < 0.05 and |average log2 FC| > 0.2 are colored by the gate in which they are upregulated (in-contact, blue; singlet, gold). (B) Top pathways in in-contact T cells from gene set enrichment analysis (GSEA), ordered by enrichment score (blue, positive; gold, negative), with log-transformed adjusted p-value shown on x-axis. (C) Alluvial plot showing min-max normalized TPM (transcripts per million) of the top 10 upregulated DEGs (ranked by log2FC) in in-contact T cells queried from the DICE (Database of Immune Cell Expression, Expression quantitative trait loci (eQTLs) and Epigenomics) database. Gene expression values are scaled between 0 and 1. Cell types on the x-axis are grouped by major immune lineage (Monocyte lineages, T cell lineages, and others) and ordered by cumulative scaled expression scores. (D) Dot plot of expression level of six genes across three image-gated categories and dual-positive (DP) T cell-monocyte droplets. (E) Heatmap displaying z-score scaled expression levels of 90 significantly upregulated DEGs in in-contact T cells. Rows (genes) are divided into three hierarchical clusters using unsupervised Ward.D2 clustering and annotated on the left based on the T cell-monocyte intermixing model: monocyte (red) and T-cell endogenous (turquoise). For visualization, columns represent single cells from randomly sampled 4,000 cells. (F) Top gene ontology (GO) pathways significantly associated with the three gene groups from panel (E), colored by combined score from enrichR, with log-transformed adjusted p-value shown on x-axis. (G) UMAP representation of T cells and manual cluster annotation. (H) Dot plot showing top markers genes for T cell clusters. (I) Meta-cluster composition differences between in-contact (blue) and singlet (gold) T cells paired by sample. (J) Boxplots of per-participant average UCell module scores for the three gene groups from panel (E) across T cell meta-clusters. For I and J, paired per-participant points are shown for singlet (gold) and in-contact (blue) gates, with grey lines connecting matched participants; boxplots show the median and interquartile range (IQR). Gates were compared within each subset by paired two-sided Wilcoxon signed-rank test with BH adjustment (adjusted p-values above brackets). For D and H, color intensity represents scaled average RNA expression levels; dot size corresponds to the percentage of cells within a given group expressing each gene. Data were derived from 12,487 in-contact, 9,272 singlet T cells and 2,455 DP droplets, from n=20 PBMC samples of patients with ATB collected at diagnosis.

To rigorously test this, we simulated an *in silico* T cell-monocyte intermixing model in which T cells transcriptomes were contaminated with incremental proportions of monocyte-derived reads (simulated T cells; ranging from 0% for pure singlets to 100% for pure doublets) and compared the resulting profiles to actual in-contact T cells (**Figure S3A**). Simulated T cell transcriptomes were most similar to in-contact T cells at a monocyte contribution of only 0.41% (designated as T-M_0.41%_), yielding a total of 139 DE genes; in contrast, the 100% doublet simulation diverged substantially from in-contact T cells, yielding 1,842 DE genes (**Figure S3A**). To determine which of the 90 genes upregulated in in-contact T cells relative to singlets (T.90 signature, **Figure 3A**) could be explained by this minor monocyte intermixing, we performed DE analysis between the simulated T-M_0.41%_ cells and actual in-contact T cells. Of the 90 genes within the T.90 signature, 12 showed no significant difference compared to the T-M_0.41%_ simulation, including top DE genes *S100A8*, *S100A9* and *LYZ* (**Figure S3B**). One additional gene (*CST3*) was expressed at significantly higher levels in the T-M_0.41%_ simulation (**Figure S3B**). Thus, the elevated expression of these 13 genes in in-contact T cells relative to singlets can be explained by a minor transcriptomic admixture with monocytes. Conversely, the remaining 77 genes from the T.90 in-contact signature were expressed at significantly higher levels in in-contact T cells than in the T-M_0.41%_ simulation, indicating that their upregulation reflects endogenous T cell activation.

The 13 monocyte-associated transcripts within the T.90 signature were co-expressed within a subset of individual cells rather than randomly distributed within all cells (**Figure S3C**). Furthermore, because both the in-contact and singlet libraries contained T cells and monocytes, we ruled out ambient RNA background as their source. Additionally, their raw UMI count distributions remained well below levels observed in DP T cell-monocyte droplets (**Figure S3D**), excluding the possibility that the monocyte-like T cells represent droplets containing an intact T cell and an intact monocyte. Taken together, these results demonstrate that the presence of monocyte-specific transcripts in in-contact T cells reflects single T cells that have acquired a small fraction of monocyte-derived RNA – most likely from localized material transfer during physical cell-cell contact.

### Metabolism, inflammation and cytotoxic signatures induced by physical contact with monocytes are universally shared across T cell subtypes

Hierarchical clustering of the T.90 signature within in-contact T cells revealed three distinct co-expression modules: Group 1 (63 genes), Group 2 (8 genes), and Group 3 (19 genes, **Figure 3E**). The 13 monocyte-associated transcripts were concentrated in Group 2 (6 genes, 75% of cluster) and Group 3 (5 genes, 26% of cluster) (**Figure 3E**). Gene ontology (GO) biological process enrichment analysis highlighted clear functional divergence among these modules: Group 1 was associated with translation and oxidative phosphorylation, Group 2 with inflammatory responses, and Group 3 with cytolytic activity (**Figure 3F**). Thus, the transcriptional signature of in-contact T cells partitions into three distinct biological axes: metabolism, inflammatory and cytotoxicity.

To determine whether this response was restricted to specific T cell lineages, we mapped these functional modules across high-resolution T cell subpopulations. Unsupervised graph-based clustering identified 14 distinct transcriptomic clusters, which were consolidated into 9 meta-clusters: CD4^+^ naïve, CD4^+^ memory, CD4^+^ Tregs, CD8^+^ naïve, CD8^+^ cytotoxic, MAIT, iNKT, γδ T cells and double-negative (CD4^-^CD8^-^) T cells (**Figures 3G**, **3H**, **Table S2**). Meta-cluster frequencies remained consistent across individual patients (**Figure S3E**). While in-contact T cells did not display a marked skewing toward any single meta-cluster (**Figure 3I**), they exhibited a statistically significant enrichment for the CD8^+^ cytotoxic KIR/KLR^+^ cluster 5 (**Figure S3F**). Conversely, singlet T cells were preferentially enriched for naïve CD8 T cells (**Figure 3I**, **S3F**).

Remarkably, when we quantified module-specific expression scores per cluster, the inflammatory, cytotoxic, and metabolic gene modules were consistently upregulated in in-contact T cells relative to singlet counterparts across all defined subsets, whether analyzed by broad meta-cluster (**Figure 3J**) or high-resolution subcluster (**Figure S3G**). These data demonstrate that physical interaction with monocytes is associated with a universal transcriptomic signature characterized by heightened activation, metabolic and effector state shared across all T cell lineages.

### The global TCR repertoire architecture is conserved between in-contact and singlet T cells

To assess whether T cell-monocyte interactions correlate with specific clonal features, we analyzed the TCR repertoire of in-contact and singlet T cells. In-contact T cells exhibited an increased frequency of expanded clonotypes (>1 cell, **Figure 4A**). However, mean clone size and global diversity indices were equivalent between groups (**Figures 4A, 4B**), indicating that the in-contact clonal expansion consists of multiple moderately sized clones rather than hyper-dominant clonotypes, thereby preserving global repertoire diversity.

**Figure 4:**
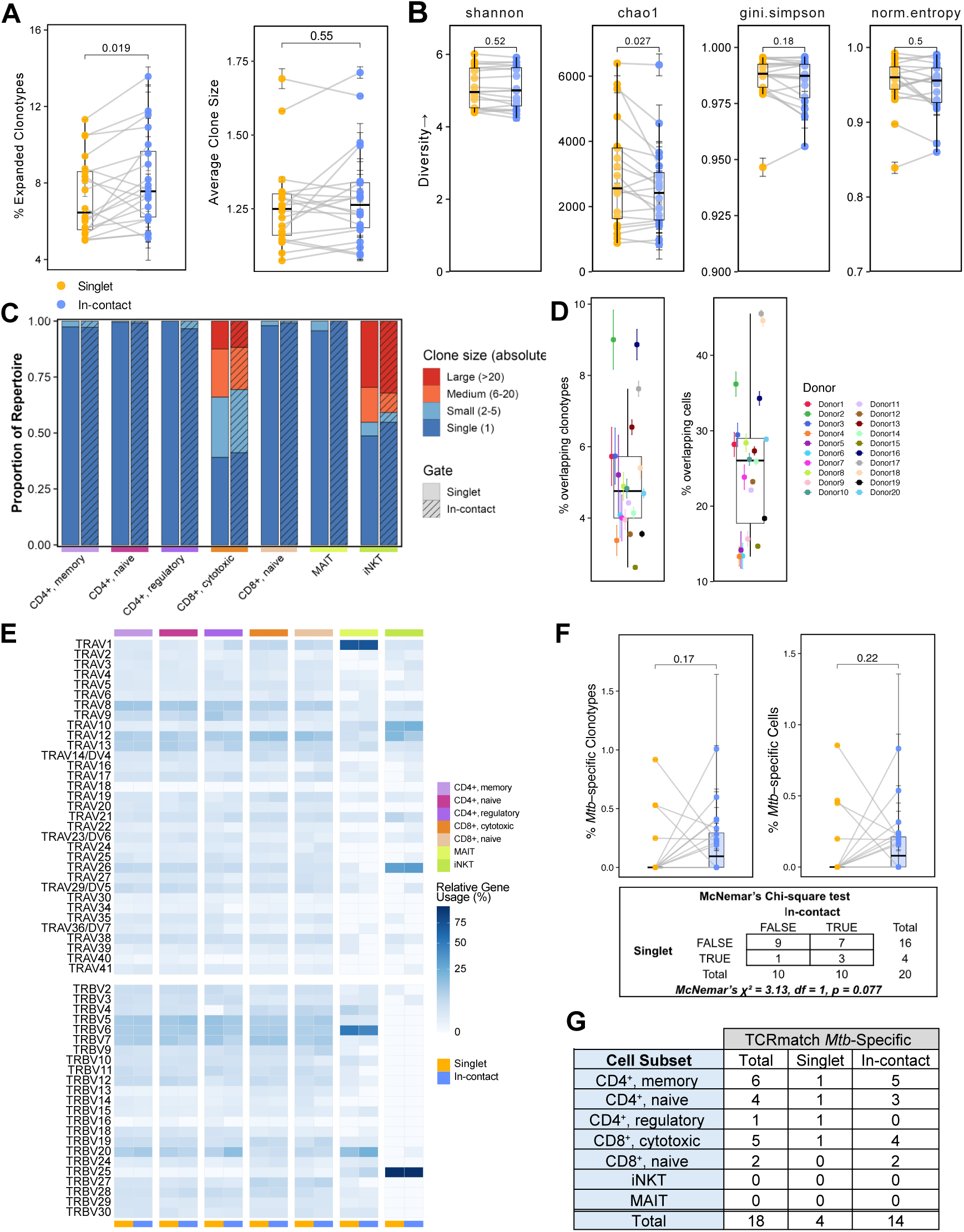
Conserved global TCR repertoire architecture and trend toward antigen-specific enrichment in in-contact T cells. (A) Clonal expansion metrics showing the percent of expanded clonotypes (with 2 or more cells per clone; left) and average clone size (right) across paired participants. (B) TCR repertoire diversity indices across four complementary metrics: Shannon index, Chao1 richness estimator, Gini-Simpson index, and normalized entropy. (C) Clonal size distribution across meta-clusters, split by in-contact (striped columns) and singlet (clear columns) T cells. Stacked bar plots display the proportion of the TCR repertoire with clonotypes categorized into four clone size bins: Single (1 cell, dark blue), Small (2-5 cells, light blue), Medium (6-20 cells, orange), and Large (>20 cells, red). Cells were downsampled to equalize total cell counts per T-cell subset within each image-gated category. (D) TCR clonotype overlap between paired singlet and in-contact T cells per participant, with the percentage of overlapping clonotypes (left) and the percentage of cells expressing a shared clonotype (right). Singlet and in-contact T cell populations for each participant were downsampled to the same number (100 iterations; points and error bars indicate mean ± SD across iterations); percentages were computed per iteration relative to the combined clonotype or cell counts for both populations for one participant. (E) Heatmap of TRAV (top) and TRBV (bottom) gene usage across meta-clusters, comparing singlet (gold) versus in-contact (blue) T cells. Color intensity was computed as the percentage of cells expressing that gene among all cells with both α and β chain sequenced in that group. (F) Proportions of Mtb-specific CDR3β clonotypes relative to total CDR3β clonotypes (left) and Mtb-specific cells relative to all cells expressing a CDR3β (right). Mtb antigen specificity was determined using TCRmatch (score threshold 0.97) against the IEDB database. The bottom panel shows a paired contingency table of number of participants with the presence (TRUE) or absence (FALSE) of Mtb-specific clonotypes in the singlet versus in-contact T cells, tested using McNemar’s chi-square test. (G) Summary table of Mtb-specific cell counts across meta-clusters split by singlet and in-contact T cells. For A, B, D and F, metrics represent participant-level mean values calculated from 100 bootstrap downsampling iterations. Boxplots display the overall median and IQR, vertical error bars represent mean ± SD across bootstrap iterations, and gray lines connect matched participant samples between singlet (gold) and in-contact (blue) T cells. Paired Wilcoxon signed-rank tests were used for statistical analysis. For all the panels, data are derived from 14,796 T cells with both TCRA and TCRB sequences detected (i.e., clusters with >50% positive cells) from n=20 PBMC samples of patients with ATB collected at diagnosis.

Subset-specific clonal analysis (considering meta-clusters in which >50% of cells had both TCRα and TCR β detected, **Figure S4A**) revealed that clonality was concentrated within specific effector lineages (CD8^+^ cytotoxic, iNKT and MAIT cells), with similar proportions of medium and large clones between in-contact and singlet T cells (**Figure 4C**). Intra-participant repertoire overlaps between in-contact and singlet T cells varied substantially: some participants displayed a highly shared repertoire (up to 9.0% overlapping clonotypes and 45.5% overlapping cells) whereas others showed minimal overlap (as low as 2.6% overlapping clonotypes and 13.3% overlapping cells; **Figure 4D**). Additionally, variable gene usage for both TCRα (TRAV) and β (TRBV) chains showed no significant differences between in-contact and singlet T cells (**Figure 4E**). Collectively, these data indicate that physical engagement with monocytes is not associated with a systematic bias in global TCR repertoire structure.

### Antigen-specific T cells trend toward enrichment within the monocyte-bound population

To determine whether physical contact with monocytes is biased toward antigen-specific T cells, we mapped the TCR sequences of in-contact and singlet T cells against public TCR databases. Initial analysis using a compiled meta-resource of *Mycobacterium tuberculosis* (Mtb)-reactive TCRs and Mtb metaclones (14) revealed comparable numbers of Mtb-reactive clonotypes and cells between in-contact and singlet T cells (**Figures S4B**, **S4C**). We next leveraged TCRmatch (15), applying a more stringent curation that restricted matches exclusively to validated Mtb epitope pairs with no documented cross-reactivity to non-Mtb pathogens (tmthus defined as not only Mtb-reactive, but Mtb-specific). Here, we observed an enrichment trend in both the frequency of Mtb-specific clonotypes and cells within the in-contact population (**Figure 4F**). Specifically, Mtb-specific TCRs were exclusively detected in in-contact T cells in 7 of 20 participants, compared to just 1 participant in singlet T cells (**Figure 4F**; McNemar’s Chi-square test, p = 0.077).

Partitioning these TCRmatch antigen-specific hits (n=18 cells total) by transcriptomic phenotype revealed that Mtb-specific T cells mapped predominantly to the CD4+ memory and CD8+ cytotoxic meta-clusters (**Figure 4G**), aligning with expected polarization of Mtb-specific T cell responses in ATB (Th1, Th1* and cytotoxic phenotypes (16,17)). Strikingly, in these two meta-clusters, Mtb-specific hits were almost exclusively restricted to the in-contact population, with only a single Mtb-specific cell detected per subtype in the singlet compartment (**Figure 4G**). Taken together, our TCR specificity analysis demonstrates that antigen-specific T cells are preferentially enriched within the monocyte-bound T cell population.

### In-contact monocytes express a universally shared metabolic signature

Next, we pivoted our analysis to the monocyte compartment. Unsupervised graph-based clustering identified 8 distinct monocyte clusters, two of which exhibited atypical transcriptomic profiles (**Figure 5A**, **Table S3**). Cluster 4 represented a low-quality subset characterized by lower-than average number of genes and transcripts per cell, an increased percentage of mitochondria genes, and a reduced percentage of ribosomal genes (**Figure S5A**). Cluster 7 formed an isolated island on the UMAP with distinct transcriptomic features (**Figure 5A**), marked by robust expression of T cell-specific marker genes (**Table S3**). Notably, this expression profile differed from the low-level monocyte gene traces observed in in-contact T cells, which did not form an independent cluster (**Figure 3G**). Furthermore, T cell marker expression levels in Cluster 7 were comparable to those of DP T cell-monocyte droplets identified during our initial filtering pipeline (**Figure S5B**). We therefore concluded that Cluster 7 represented intra-sample T cell-monocyte doublets that bypassed initial exclusion; consequently, both Clusters 4 and 7 were removed to ensure downstream analyses focused strictly on high-quality single cells.

**Figure 5:**
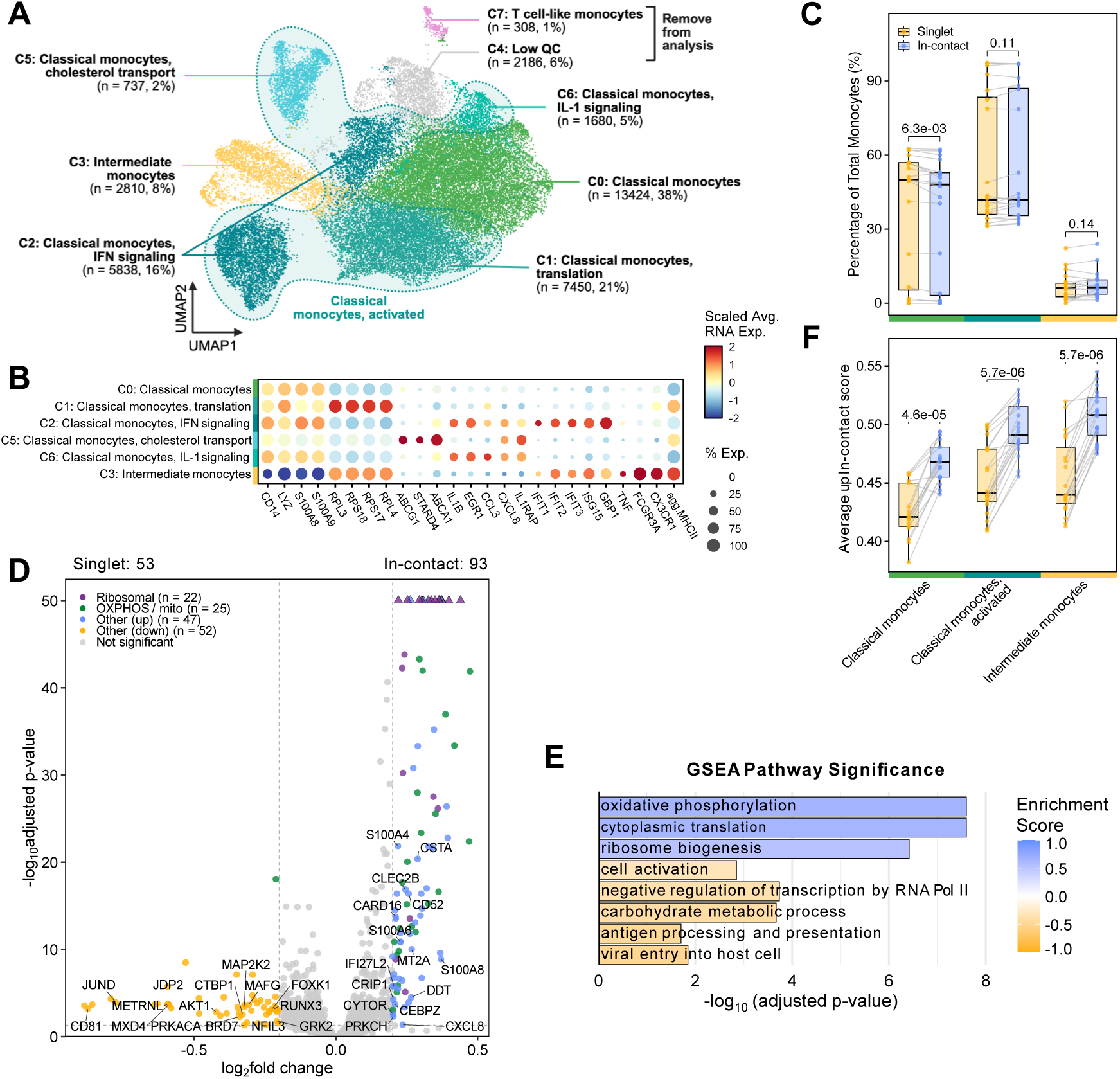
In-contact monocytes are associated with specific gene expression signatures. (A) UMAP representation of monocytes and manual cluster annotation. (B) Dot plot showing top markers genes for monocyte clusters. Color intensity represents scaled average RNA expression levels; dot size corresponds to the percentage of cells within a given cluster expressing each gene. (C) Meta-cluster composition differences between in-contact (blue) and singlet (gold) monocytes paired by sample, using non-parametric paired Wilcoxon tests. (D) Volcano plot of DEGs comparing in-contact versus singlet monocytes. Genes with BH adjusted p-value < 0.05 and |average log2 FC| > 0.2 are color-coded: ribosomal genes (purple), OXPHOS/mitochondrial genes (green), other DEGs upregulated in in-contact monocytes (blue), other DEGs upregulated in singlet monocytes (gold), and non-significant genes (gray). Triangles represent highly significant genes capped at -log_10_ BH-adjusted p-value > 50. (E) Top pathways in in-contact monocytes from GSEA analysis, ordered by enrichment score (blue, positive; gold, negative), with log-transformed adjusted p-value shown on x-axis. (F) Boxplots of per-participant average UCell module scores of 93 DEGs significantly enriched in in-contact monocytes across meta-clusters. For C and F, paired per-participant points are shown for singlet (gold) and in-contact (blue) monocytes, with grey lines connecting matched participants; boxplots show the median and IQR. Singlet and in-contact monocyte populations were compared using a paired two-sided Wilcoxon signed-rank test with BH adjustment (adjusted p-values above brackets). Data were derived from 11,915 in-contact and 10,448 singlet monocytes, from n=20 PBMC samples of patients with ATB collected at diagnosis.

Based on top marker gene expression (**Figure 5B**, **Table S3**), the remaining clusters were categorized into intermediate monocytes (Cluster 3), classical monocytes (Cluster 0) and specialized classical monocytes displaying distinct activation signatures: interferon signaling (Cluster 2), IL-1 signaling (Cluster 6), cholesterol metabolism (Cluster 5), and translation (Cluster 1). These activated subsets were consolidated into an “activated classical monocyte” meta-cluster. Meta-cluster composition was highly consistent across participants (**Figure S5C**). Overall, in-contact and singlet monocytes exhibited minimal differences in meta-cluster and individual cluster frequencies, with classical monocytes slightly enriched among singlets (**Figures 5C**, **S5D**).

Despite equivalent subset proportions, differential expression analysis revealed transcriptomic divergence, identifying 93 upregulated and 53 downregulated genes in in-contact monocytes relative to singlets (**Figure 5D**, **Table S4**). Upregulated genes were strongly associated with translation (marked by ribosomal genes), and, mirroring our observations in T cells, oxidative phosphorylation (including multiple *COX* and *ATP* family genes; **Figures 5D, 5E**). Like the T cell compartment, this in-contact signature was not confined to a specific monocyte lineage but was universally upregulated across all meta-clusters (**Figure 5F**), and individual clusters (**Figure S5E**). Together, these findings demonstrate that CD14^+^ monocytes physically interacting with T cells share a universal, highly active metabolic state defined by elevated oxidative phosphorylation and translational activity.

### Partner-derived transcriptomic traces are unidirectional from monocytes to T cells

Having identified clear monocyte-derived gene traces within in-contact T cells, we next investigated whether the reciprocal phenomenon could be observed within the monocyte compartment. In contrast to T cells, our DE analysis of in-contact and singlet monocytes revealed no T cell-specific transcripts (**Figure 5D**), suggesting an absence of reciprocal material transfer. Relaxing the minimum percent-expression cutoff from 25% to 0% in the DE analysis yielded 98 additional genes upregulated in in-contact monocytes (**Figure S5F**). Among these 98 genes, 57 were more highly expressed in T cells than in monocytes according to the DICE database (18) (**Figure S5G**). Both the percentage of expressing cells and expression level of these 57 T cell-associated genes were extremely low in in-contact monocytes – virtually indistinguishable from background singlet monocytes and far below those of DP T cell-monocyte droplets (**Figure S5H**). Furthermore, these T cell transcripts were seldom co-expressed in individual monocytes (**Figure S5I**). This observation contrasted sharply with the robust monocyte gene signature in in-contact T cells, which clearly separated them from singlets and was broadly shared across individual T cells (**Figure 3D**, **S3C**, **S3D**). Ultimately, these findings demonstrate that partner-derived transcriptomic traces are unidirectional, occurring predominantly from monocytes to T cells.

### Coincidental doublets express intermediate cell-cell interaction signatures

Having characterized in-contact T cells and monocytes relative to their singlet counterparts, we next interrogated the transcriptomic profiles of coincidental doublets. Under the hypothesis that coincidental doublets represent purely random spatial co-encapsulations, their transcriptomes were expected to mirror those of singlets. However, the in-contact DE signature of T cells (T.90 signature, **Figure 3A**) and monocytes (M.93 signature, **Figure 5D**) exhibited an intermediate level of expression in coincidental T cells and monocytes – higher than in singlets, but significantly lower than in in-contact doublets (**Figure 6A**). This indicated that coincidental doublets are not transcriptomically identical to singlets and instead share features with in-contact populations. This intermediate state was observed across both T cell (**Figure S6A**) and monocyte meta-clusters (**Figure S6B**). Quantifying the top 10% highest-scoring cells within each category confirmed that coincidental T cells harbor the ‘in-contact signature’ at a significantly higher frequency than singlet T cells, but at a lower frequency than in-contact T cells (**Figure 6B**). Monocytes exhibited greater inter-individual variability; while the majority of participants displayed a trend similar to T cells, the difference did not reach statistical significance (**Figure 6B**). Least-squares mixture model of the coincidental distribution showed that the expression profile of coincidental cells is consistent with a mixture of about 1/3 singlets (35-40%) and 2/3 (60-65%) in-contact cells for both T cells and monocytes (**Figure 6C**). Together, these findings demonstrate that coincidental doublets are not entirely random; rather, a significant fraction display residual contact signatures, likely representing in-contact pairs that dissociated immediately prior to laser beam interrogation. Furthermore, these data indicate that transcriptomic interaction signatures persist temporarily after physical cell-cell separation, particularly in T cells.

**Figure 6:**
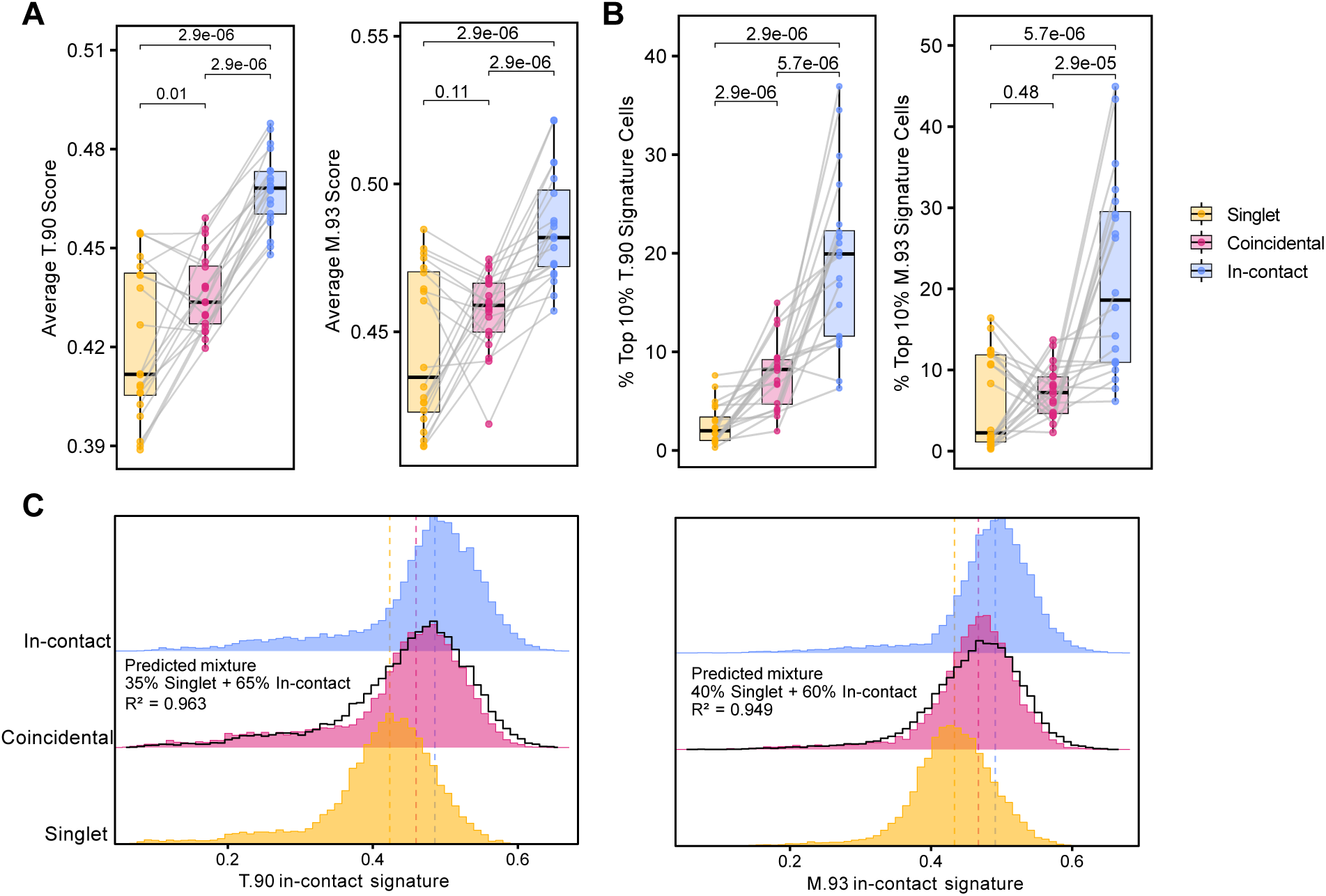
Coincidental doublets express intermediate in-contact signatures. (A) Boxplots of per-participant average UCell module scores for 90 DEGs found up in in-contact T cells compared to singlet T cells (T.90 signature; left), and 93 DEGs found up in in-contact monocytes compared to singlet monocytes (M.93 signature; right). Both signature scores are calculated using UCell. (B) Boxplots of per-participant percentage of cells within the top 10% for the T.90 signature (left), and M.93 signature (right). (C) Ridge plot showing the distribution of UCell module scores for T.90 signature (left) and M.93 signature (right) across singlet (gold), coincidental (pink), and in-contact (blue) gates. Vertical dashed lines indicate median score values for each category. The black stepped line overlaid on the coincidental ridge represents the predicted mixture distribution modeled via least-squares optimization as a linear combination of singlet and in-contact density profiles. The resulting predicted mixture proportions and the goodness-of-fit statistic are showed next to the stepped line. For A and B, paired per-participant points are shown for singlet (gold), coincidental (pink) and in-contact (blue) gates, with grey lines connecting matched participants; boxplots show the median and IQR. Gates were compared within each subset by paired two-sided Wilcoxon signed-rank test with BH adjustment (adjusted p-values above brackets). Data were derived from 9,272 singlet, 12,022 coincidental and 12,487 in-contact T cells; 10,448 singlet, 11,002 coincidental and 11,915 in-contact monocytes from n=20 PBMC samples of patients with ATB collected at diagnosis.

### Dual-lineage droplet analysis validates paired T cell-monocyte interaction signatures

Finally, we evaluated the transcriptomes of dual-lineage T cell-monocyte droplets (i.e., droplets harboring both T cell and monocyte gene signatures at high expression levels), which correspond to intra-sample T cell-monocyte doublets. While present across all sorted libraries as technical artifacts, dual-lineage droplets in the in-contact condition also contain a bona fide biological fraction (∼3.4% sorted T cell-monocyte doublets remaining undissociated post-sort; **Figure 1D**). Consequently, comparing dual-lineage droplets from singlet versus in-contact libraries provides additional insight into intact T cell-monocyte interaction profiles.

Dual-lineage droplets were defined by co-positive T cell and monocyte module scores (DP droplets, **Figure 2C**), or assignment to the T cell-like monocyte cluster (Cluster 7, **Figure 5A**). As expected, dual-lineage droplets were significantly enriched in the in-contact population compared to singlets (**Figure S7A**). DE analysis between in-contact and singlet dual-lineage droplets yielded few DE genes (*MT-ND3*, *ZEB1*, *TOMM7* upregulated; *HDAC9*, *JUND*, *METRNL* downregulated; **Figure S7B**), and GSEA revealed enrichment of translation-related pathways in the in-contact category (**Figure S7C**). Crucially, the in-contact T cell and monocyte signatures identified in dissociated cells (T.90 signature, **Figures 3A**; M.93 signature, **Figure 5D**) were markedly elevated in in-contact dual-lineage droplets (**Figure S7D**). Restricting the analysis to genes not overlapping between the two signatures, the T.90 and M.93 signature scores displayed a stronger intracellular correlation within in-contact than singlet dual-lineage droplets (median Spearman ρ ∼ 0.65 versus ∼ 0.47; paired *p* = 0.049; **Figure S7E**). This confirms that in-contact signatures are co-enriched within intact doublets as paired interaction profiles.

UMAP clustering of dual-lineage droplets resolved three subpopulations: a primary TCR signaling cluster (Cluster 0), a Wnt/Ca^2+^ signaling cluster (Cluster 1), and a minor platelet-associated cluster (Cluster 2), consistently represented across participants (**Figures S7F, S7G, S7H**). Dual-lineage droplets were preferentially enriched for the TCR signaling cluster over the Wnt signaling cluster in the in-contact group relative to singlets, while the platelet-associated cluster was uniquely present in the in-contact group (**Figure S7I**). Together, these results confirm that in-contact signatures derived from dissociated single cells reflect the transcriptomic state of intact, non-dissociated biological doublets, while revealing additional functional features of T cell-monocyte interactions, including localized TCR engagement and minor platelet involvement.

## Discussion

In this study, we present the first high-resolution single-cell transcriptomic characterization of physically interacting T cell-monocyte doublets isolated using real-time spectral imaging cell sorting. By integrating high-throughput morphometric and spatial fluorescence metrics from the BD FACSDiscover™ S8 platform into our physical cell deconvolution workflow, we successfully switched from traditional computational inference of doublets to direct visual verification of physical cell-cell engagement. Unlike conventional flow cytometry, which infers interactions based solely on co-expression of surface markers like CD3 and CD14, real-time imaging provides gating features that can distinguish true biological complexes from coincidental artifacts. This distinction is critical: by purging non-interacting proximal cells lacking synaptic polarization, we uncovered T cell and monocyte transcriptomic signatures of true physical interactions – such as universal oxidative phosphorylation programs and unidirectional RNA transfer – that were masked by technical noise in previous analyses.

A central strength of our experimental design is the rigorous, multi-layered pipeline implemented to validate single-cell transcriptomes following fluidic dissociation. Because droplet-based scRNA-seq inherently generates random co-encapsulations, and a small fraction of physical doublets escape sorting-induced dissociation (∼3.4%, **Figure 1D**) entering the microfluidic channel intact, purging artifactual and non-dissociated doublets was essential. This was particularly critical for evaluating intercellular material transfer; confirming that single cells were truly dissociated ensured that partner-derived transcripts did not simply represent reads from non-dissociated physical doublets.

Crucially, we identified a robust panel of monocyte-specific transcripts – including top DE genes *S100A8*, *S100A9,* and *LYZ* – that were strikingly enriched within dissociated, previously in-contact T cells. By deploying *in silico* doublet modeling, we found that as little as 0.41% monocyte read intermixing was sufficient to reproduce this signature without invoking T cell-intrinsic expression. Furthermore, these monocyte-derived transcripts were co-expressed within individual T cells at quantitative levels far below those of intact doublets, confirming they do not represent doublet contamination. Ambient RNA contamination was similarly excluded, as both singlet and in-contact libraries contained co-isolated T cells and monocytes and were thus exposed to identical backgrounds of ambient RNA. Instead, these findings point to localized leakage of monocyte-derived genetic material occurring during physical cell-cell engagement. Interestingly, this transcript acquisition was strictly unidirectional: in-contact monocytes contained no reciprocal T cell-specific gene traces. Intercellular RNA exchange across the immunological synapse is well-documented, occurring via microvesicles (19), exosomes (20), or tunneling nanotubes during direct cell-cell contact (21). Although the exact mechanism by which circulating T cells acquire monocyte transcripts remains to be fully elucidated, this finding underscores the advantage of physical deconvolution over purely bioinformatic deconvolution strategies (e.g., PIC-seq) for doublet analysis. Computational deconvolution algorithms would inevitably classify such unidirectional RNA transfer as a double-cell artifact or ambient noise, thereby completely missing bona fide intercellular RNA exchange. Finally, this highlights the superior resolution afforded by imaging flow cytometry for doublet isolation: while non-imaging physical deconvolution previously enabled us to detect dual-lineage gene expression in in-contact T cells and monocytes (8), the resolution was insufficient to determine that this transcript transfer was strictly unidirectional from monocytes to T cells and affected all sub-lineages within T cells.

Beyond the acquisition of monocyte-derived transcripts, in-contact T cells exhibited a marked endogenous response characterized by elevated cytotoxicity and metabolic activation. While the upregulation of cytotoxic pathways aligns with established effector T cell functions and prior T cell-monocyte doublet analyses (4,6,8), the robust metabolic signature – defined by the extensive co-upregulation of oxidative phosphorylation (OXPHOS) machinery (including mitochondrial, COX, and ATP gene families) alongside ribosomal genes – represents a novel insight obscured in prior doublet studies. This suggests that image-guided sorting selectively captures T cells actively or very recently engaged in immunological communication, effectively preserving transient, contact-dependent metabolic states that are typically lost in bulk or non-imaging single-cell protocols.

When evaluating the TCR repertoire, we observed a significantly higher frequency of expanded clonotypes in in-contact T cells compared to singlets. However, overall diversity metrics and mean clone sizes remained equivalent between the two populations, indicating that clonal expansion within T cell-monocyte complexes comprises multiple moderately expanded clones rather than a few hyper-dominant clonotypes. Using TCRmatch to query validated Mtb TCRβ sequences, we detected a trend toward enriched Mtb-specific TCRs in the in-contact T cell pool (i.e., presence in a greater number of participants compared to singlets). Notably, these matched Mtb-specific TCRs mapped almost exclusively to CD4^+^ memory and CD8^+^ cytotoxic transcriptomic meta-clusters. In ATB, Mtb-reactive T cells predominantly partition into Th1/Th1* memory and cytotoxic effector states (16,17), directly matching the phenotypic signature of our mapped hits. The lack of statistical significance in overall Mtb-specific clonotype frequencies likely reflects the inherent limitations of public TCR repositories, which capture only a small, non-random fraction of total pathogen-reactive repertoires. For Mtb, the IEDB is heavily biased towards well-defined immunodominant antigens, thus our analysis preferentially detects these well-documented epitopes rather than sampling the broader repertoire evenly. Furthermore, the small number of Mtb-specific cells detected over the entire cohort (n=18) limits statistical power. Future studies leveraging matched, participant-specific antigen-specific TCR reference sets will be vital to fully resolve the antigen-specific landscape within circulating T cell-monocyte complexes.

In the monocyte compartment, in-contact cells similarly displayed a universal metabolic activation state marked by elevated oxidative phosphorylation and translation across all subclusters, mirroring our findings on T cells. Interestingly, a predominant reliance on OXPHOS machinery in immune cells reflects an energetic state aligned with long-term survival or differentiation rather than rapid proliferation – which relies primarily on aerobic glycolysis (22). This suggests that circulating T cell-monocyte complexes do not represent transient, highly proliferative inflammatory blasts, but rather energetically optimized cellular interaction units poised for sustained effector function, and efficient translational output.

Interestingly, we did not observe the prominent upregulation of MHC-II antigen-presentation genes that dominated our previous non-imaging physical deconvolution study (8). This discrepancy is most plausibly explained by the enhanced resolution and purity offered by image-guided spectral sorting. Traditional non-imaging sorting relies strictly on two-dimensional scatter parameters (e.g., FSC/SSC height vs. area), which can inadvertently capture non-specific multi-cellular aggregates as doublets. In ATB, intermediate CD14^+^CD16^+^ monocytes are preferentially expanded and activated, and express high levels of MHC-II (23); their heightened activation state likely increases their cell-adhesion properties and propensity to form stochastic, multi-cellular clusters during sample preparation. In conventional scatter-based sorting, these aggregates would be artificially over-represented, skewing the overall in-contact monocyte population towards an MHC-II gene signature. By enforcing strict visual confirmation of true 1:1 cellular interaction units, image-guided sorting effectively excludes these non-specific aggregates. Consequently, the transcriptomic data from the current study reflect the true, unconfounded gene expression profile of monocytes physically engaged with T cells, which is defined by metabolic signatures rather than MHC-II transcriptional induction.

Findings derived from T cell and monocyte in-contact signatures may carry direct implications for TB pathogenesis. Monocytes are a principal cellular reservoir for *Mtb* in vivo, trafficking into the lung where they differentiate into macrophages within the granuloma, a relationship in which T cell-macrophage communication is increasingly recognized as central to disease outcome, through distinct types of contact-dependent signals that recruit, sustain, or activate macrophages to kill ingested bacteria (24). Similar functional consequences are observed in blood: imaging-based analysis of *ex vivo* T cell-monocyte conjugate formation found that, despite reduced conjugate formation in active disease, T cells that did engage monocytes were hyper-responsive (25), suggesting these interactions remain functional, not only incidental, in the periphery. Our data raise the possibility that such contacts serve dual, non-mutually exclusive roles: initiating antimicrobial programs, as reflected by the antigen-specific enrichment trend and heightened metabolic activity we observed, or terminating them, as suggested by the concurrent cytotoxic signature and enrichment of CD8+ cytotoxic subsets, potentially reflecting elimination of infected or antigen-presenting monocytes. Because complex frequencies fluctuate with disease state and anti-TB therapy (2), applying our framework to a longitudinal cohort could clarify whether this population marks active antimicrobial engagement, its resolution, or both.

An unexpected finding was the distinct transcriptomic profile of coincidental doublets. Rather than mirroring singlet controls – as anticipated for purely random fluidic co-encapsulations – coincidental doublets displayed an intermediate transcriptional state, harboring in-contact activation signatures at a lower magnitude across both T cell and monocyte subsets. Mathematically, this intermediate population was best modeled as a mixture of 35-40% singlets and 60-65% in-contact T cell or monocytes. This indicates that coincidental gates are not purely stochastic, but instead capture a substantial subpopulation of cells carrying a partial interaction signature. We propose two, non-mutually exclusive, explanations for this phenomenon. First, some coincidental events may represent genuine physical doublets that were mechanically disrupted by high fluidic pressure and velocity immediately prior to laser interrogation, such that the resulting single cells retain a residual transcriptomic “imprint” of their prior engagement. Second, and equally plausible, some coincidental events may instead capture cell pairs caught in the earlier stages of synapse formation – proximal and beginning to signal, but not yet compacted into the tight, spatially correlated configuration required for classification as in-contact. Because our imaging measurement provides only a single snapshot in time, our data cannot distinguish a population caught while disengaging from one caught while engaging, and both processes likely contribute to the coincidental gate to some degree. Regardless of directionality, this finding indicates that cell-cell contact leaves a detectable transcriptomic signature across a continuum of physical engagement states, and that true singlet T cells and monocytes – rather than coincidental doublets – remain the most rigorous control population for comparisons against in-contact cells.

Finally, our evaluation of dual-lineage doublets validated the association between the dissociated T cell and monocyte in-contact signatures. The T.90 and M.93 signatures exhibited remarkable co-enrichment (Spearman ρ ∼ 0.65) within intact doublet transcriptomes. Although these in-contact signatures were universally shared across T cell and monocyte subtypes in our cohort, this combined analytical framework – integrating both dissociated partners and intact complexes – provides a powerful template for mapping sub-lineage specific interaction signatures in other doublet populations or disease contexts displaying greater heterogeneity. Furthermore, profiling intact complexes revealed additional features of active physical engagement that were absent in dissociated cells – most notably an enrichment for a TCR signaling cluster. The selective presence of this cluster in intact doublets suggests that uncoupling triggers rapid downregulation or decay of TCR-signaling mediated transcripts. Lastly, we also identified a distinct subset of non-dissociated doublets characterized by high expression of platelet-associated transcripts (e.g., *PPBP*, *PF4*). This indicates that platelets may be responsible for increased adherence in some T cell-monocyte interactions, though such complexes represented only a minor fraction of the total intact doublet pool.

Our study has several limitations. First, our cohort was restricted to patients with ATB. Evaluating healthy control cohorts alongside disease states will be essential to determine whether the universal in-contact signatures we observed in both T cells and monocytes reflect ATB-driven increases in cell adhesiveness or represent a fundamental, steady-state property of circulating T cell-monocyte doublets. Second, our image-guided analysis categorized doublets primarily into synaptic versus coincidental configurations; however, richer morphological heterogeneity exists within these gates. Future studies incorporating additional spatial parameters – such as localized CD3 intensity, receptor clustering, or contact-surface area – may provide further resolution to isolate recently conjugated, functionally active pairs. Finally, while our image-guided sorting isolates true 1:1 contact pairs, the physical dissociation step relies on mechanical stressors in the cell sorting process. A complex’s susceptibility to dissociation *in vitro* may be governed by synaptic maturation or adhesion molecule density, meaning non-dissociated doublets might over-represent high-affinity interactions and consequently be depleted in the dissociated single-cell pool. Although we mitigated this bias by incorporating the analysis of dual-lineage droplets – containing non-dissociated T cell-monocyte doublets – in our workflow (**Figure S7**), they represented a very small fraction of the overall dataset. Future studies leveraging this shear-resistance feature could selectively isolate these high-affinity doublet populations to uncover the transcriptomic determinants of synaptic stability.

Overall, this study establishes that combining real-time spectral image-guided sorting with physical deconvolution yields an unprecedented, high-fidelity platform for decoding human cell-cell interactions. By rigorously distinguishing true physically interacting pairs from coincidental artifacts, we revealed universal metabolic and cytotoxic programs orchestrating T cell-monocyte communication in active TB, uncovered evidence of unidirectional RNA transfer, and established a scalable blueprint to map physical immune networks across human health and disease.

## Material and methods

### Study cohort and samples

Human study participants were enrolled at the field site of the South African Tuberculosis Vaccine Initiative (SATVI; University of Cape Town, Western Cape Province, South Africa), and the Universidad Peruana Cayetano Heredia (Peru). Ethical approval to carry out this work was granted and maintained by the Institutional Review Board of the La Jolla Institute for Immunology and the Universidad Peruana Cayetano Heredia, and the Human Research Ethics Committee of the University of Cape Town, in accordance with regulations for the protection of human subjects. All clinical investigations were conducted according to the principles expressed in the Declaration of Helsinki, and all participants (or guardians for participants <18 years old) provided written, informed consent before participation in the study. ATB disease was defined by 1) the presence of clinical symptoms and/or radiological/histological evidence of pulmonary TB, and 2) microbiological confirmation via *Mtb*-specific molecular or culture testing. PBMC were isolated from whole blood samples by density gradient centrifugation using Ficoll-Paque^®^ PLUS (Cytiva) according to the manufacturer’s instructions. Isolated PBMC were resuspended at up to 10 million cells per milliliter in fetal bovine serum (FBS; Gemini Bio) containing 10% dimethyl sulfoxide (DMSO; Sigma-Aldrich) and cryopreserved in liquid nitrogen.

### PBMC thawing

Cryopreserved PBMC were rapidly thawed in a 37°C water bath for 2 minutes and immediately transferred into 9 ml of cold complete medium consisting of RPMI 1640 with L-Glutamine and 25 mM Hepes (Omega Scientific), supplemented with 5% human AB serum (GemCell), 1% penicillin/streptomycin (Gibco), 1% GlutaMAX (Gibco), and 20 U/mL Benzonase^®^ Nuclease (Millipore). Cells were centrifuged, resuspended in complete medium, and evaluated for cell concentration and viability using trypan blue exclusion and a hematocytometer. PBMC were maintained at 4°C prior to downstream antibody staining and acquisition.

### Ex vivo antibody staining

For surface immunophenotyping, up to 10 × 10^6^ thawed PBMC were resuspended in 95 μl of phosphate-buffered saline (PBS) containing 5 μl of Human BD Fc Block^TM^ (BD Biosciences) and incubated for 10 min at room temperature. All fluorochrome-conjugated primary antibodies were centrifuged at 10,000g for 5 minutes at 4°C prior to use to pellet aggregates. Predetermined optimal volumes of fluorochrome-conjugated anti-human antibodies (5 μl anti-human CD3 BB515, clone UCHT1, BD Biosciences; 4 μl anti-human CD14 PE Dazzle 594, clone HCD14, Biolegend; 2 μl anti-human CD19 APC, clone HIB19, Biolegend) and 1 μl viability dye (Fixable viability stain 440UV, BD Biosciences), along with 10 μl of Brilliant Stain Buffer (BD Biosciences), and 70 μl flow cytometry buffer (PBS containing 2 mM EDTA [pH 8.0] and 0.5% bovine serum albumin [BSA]) were added to each sample. TotalSeq™-C oligonucleotide-conjugated antibodies (Biolegend) were also added at this step at 0.01mg/mL final concentration (one distinct antibody per sample), and samples were incubated for 30 min at room temperature in the dark. Single-stained cellular controls and compensation/unmixing beads were prepared concurrently using identical staining protocol and volumes. Following incubation, cells were washed twice in flow cytometry buffer, resuspended in 0.5mL of flow cytometry buffer and stored at 4°C protected from light for up to two hours prior to acquisition.

### Image- and spectral-guided cell sorting

Samples were acquired and sorted using a BD FACSDiscover^TM^ S8 Cell Sorter with BD SpectralFX^TM^ and BD CellView^TM^ Image Technology with the standard 5-laser configuration (349 nm, 405 nm, 488 nm, 561 nm, 637 nm), 6 imaging detectors (3 scatter, 3 fluorescence PMTs) and a spectral array of 78 APD detectors, operating on BD FACSChorus^TM^ software version 5.4. The 85μm nozzle with standard settings (35 psi, 57kHz) was used, and instrument and imaging calibration QC were performed using BD FACSDiscover^TM^ Setup Beads and BD CellView^TM^ Calibration Beads, respectively. Drop delay was determined using BD FACS^TM^ Accudrop Beads. Gain settings for LightLoss (Violet) and SSC were adjusted to optimize resolution of immune cells within PBMC for the primary gate. QC-determined gain settings were used for all other detectors. An unstained sample, single-stained cell and cell mimic controls were acquired and used to optimize the unmixing matrix applied to all samples. For imaging optimization, the Region of Analysis (ROA) was adjusted based on SSC, then Pixel Threshold for each imaging detector was set appropriately, followed by adjustments to channel settings to minimize overall background. All samples were acquired and sorted at a flow rate of <8,000 events/sec. Purity mode was used to sort up to 3,000 singlet CD3^+^CD14^-^CD19^-^ T cells, singlet CD3^-^CD14^+^CD19^-^ monocytes, coincidental T cell-monocyte doublets, and in-contact T cell-monocyte doublets (gating strategy **Figure 1B, S1A**) per sample. For some samples, a small fraction of each of the sorted populations was re-acquired on the same instrument directly after sorting, to confirm that the sort resulted in the physical separation of in-contact doublets. Samples were processed into two separate experiment runs with 10 samples each. For each experiment run, all samples were sorted and pooled into three separate collection tubes: i) singlet T cell + singlet monocytes; ii) coincidental and iii) in-contact T cell-monocyte doublets, which were then prepared and sequenced as separate libraries. Fluorescence and imaging data were analyzed using FlowJo^TM^ software (v10.10). Imaging data was visualized via the BD CellView™ Lens plugin (v2.1). Statistical analyses and graphing were performed using GraphPad Prism (v10.4).

### Single-cell RNA and TCR sequencing

Single-cell RNA sequencing was performed using the droplet-based 10X Genomics platform according to the manufacturer’s instructions. Following cell sorting, cells were spun down, and single-cell libraries were prepared as per the manufacturer’s instructions (10X Genomics). Samples were processed using 10 × 5’v3 chemistry as per the manufacturer’s recommendations. The library preparation was performed using Chromium GEM-X Single Cell 5’ Kit v3 (Dual index) with feature Barcode technology kit and chromium Single Cell Human TCR Amplification Kit. Libraries were sequenced using the Illumina NovaSeq 6000 sequencing platform.

### Single-cell RNA sequencing data analysis

#### Pre-processing

FASTQ files from the single-cell libraries were processed with 10x Genomics Cell Ranger (v8.0.1) (26) using two pipelines. Gene expression and hashtag (HTO) demultiplexing libraries were processed with cellranger multi, aligning reads to the human GRCh38 reference transcriptome (2024-A; GENCODE v44/Ensembl 109). TCR repertoire libraries were processed separately with cellranger vdj pipeline, using the human GRCh38 V(D)J reference (10x reference v7.1.0). After this step, data were converted into demultiplexed outputs, and cells with zero or more than one positive sample barcode detected were discarded (**Figure 2B**). A Seurat object was built with Cell Ranger outputs using R package Seurat (v5.0.3) (27) and R (v4.3.2). During library preparation, a subset of cells remained in the supernatant for the singlet and coincidental sorted fractions. To maximize cell recovery, these supernatant cells were collected and sequenced as separate libraries, then merged back into their corresponding main libraries (singlet and coincidental, respectively) for all downstream analyses. As an initial round of quality control, we removed cells with a percentage of mitochondrial genes lower than 5% or a total number of genes per cell less than 300, resulting in 76,918 droplets.

#### Participant-specific gene aggregation

To reduce the variance incurred by the diversity and highly individual-specific expression patterns of TCR, immunoglobulin (IG), and MHC genes, which can potentially lead to the formation of participant-specific small clusters that do not represent biologically meaningful subsets, we aggregated their raw counts into broader gene family subgroups. Specifically, individual TCR genes were aggregated into four subgroups (TCRA, TCRB, TCRG, and TCRD); IG genes into three subgroups (IGH, IGL, and IGK); and MHC genes into two subgroups (MHCI and MHCII). Certain marker genes (TRAV1-2, TRAJ33, TRAJ12, TRAJ20, TRGV9, TRDV2, TRAV10, TRAJ18, and TRBV25-1) were excluded from aggregation and retained individually, as they serve as key features for annotating specific T cell clusters downstream. For all other targeted genes, the individual raw counts were removed from the count matrix, and the summed raw counts for each respective subgroup across individual cells were incorporated into the matrix instead. Gene aggregation was applied exclusively for dimensionality reduction, graph-based neighbor construction, and cell clustering to prevent participant-specific clustering artifacts. For all downstream analyses, the original unaggregated count matrices containing individual TCR, IG, and MHC genes were retained. These unaggregated matrices were subsequently normalized and variance-stabilized using the function ‘NormalizeData’ and the function ‘SCTransform’ (regressing out percent.mt) to preserve full gene-level resolution.

#### Doublet rate inference

The number of intra-sample doublets in each run was estimated from the hashtag (HTO) demultiplexing data as shown in **Figure S2G**. PBMC from each participant were labeled with a distinct hashtag and sorted cells pooled into a single tube at its own density, so the estimated number of inter-sample doublets formed by random co-encapsulation for a given pair of hashtags was determined by the relative cell densities for each participant. For each sequencing run, the proportion of cells for each participant was calculated as p(i) = n(i) / N, where n(i) is the number of cells assigned to participant i and N is the total number of cells. For a given pair of participants i and j, the expected number of inter-sample doublets is proportional to N × 2 × p(i) × p(j), with the proportionality constant being a library-specific doublet factor (DF).

To calculate the expected number of intra-sample doublets, the observed inter-sample doublet count for each hashtag pair was regressed against its inferred value N × 2 × p(i) × p(j) using a linear model through the origin; the fitted slope gave the library-specific DF, estimated separately for each loaded lane. The number of expected intra-sample doublets for each participant was then inferred as N × DF × p(i)², and summed across participants to yield the expected artificial intra-sample doublet count.

In addition to the artificial intra-sample doublets described above, an additional category of intra-sample doublets is expected to be present in the dataset. These correspond to genuine in-contact doublets that were not dissociated after bulk sort (biological intra-sample doublets, **Figure S2A**). The frequency of these non-dissociated doublets was measured directly (**Figure 1D**) and multiplied by the corresponding cell number for each participant to obtain the biological intra-sample doublet count for each gate. The total inferred intra-sample doublet count was the sum of the artificial and biological components (**Figure 2E**).

#### Doublet identification and removal

To remove intra-sample doublets, the dual expression approach and the RNA Features approach were applied sequentially (**Figure 2A**).

1. <u>Dual expression approach</u>: After TCR gene aggregation, normalization and variance stabilization were performed using the function ‘SCTransform’ (v2 workflow, setting method = “glmGamPoi”, and regressing out percent.mt). Canonical T cell and monocyte gene module scores were calculated using the normalized SCT data assay. The canonical T-cell module was defined using *CD3E*, *CD2*, *CD3D*, *CD3G*, *IL7R*, and the aggregated TCR genes (agg.TCRA and agg.TCRB), while the canonical monocyte module comprised *S100A9*, *S100A8*, *LYZ*, *CD14*, *VCAN*, *CST3*, and *TYROBP*. For each droplet, T cell and monocyte module scores were computed by taking the sum of normalized expression values across all detected genes within each defined module set. Given the expected 1:1 sorting ratio of T cells to monocytes, the median (50^th^ percentile) of the continuous score distribution was selected as the cutoff for each module score. Droplets exceeding the 50^th^ quantile for both T-cell and monocyte scores were identified as dual-positive (DP) T cell-monocyte droplets and removed. Following removal of DP droplets, the remaining droplets were re-clustered to assign each to a major lineage. Normalization and variance stabilization were repeated on this subset using the function ‘SCTransform’ (v2 workflow, regressing out percent.mt). The function ‘FindNeighbors’ (setting dims = 1:30, k.param = 50) and the function ‘FindClusters’ (setting resolution = 0.4) were performed on the SCT assay, and cells were embedded in two dimensions using the function ‘RunUMAP’ (setting dims = 1:30, metric = “euclidean”). The resulting clusters were assigned to either T cell or monocyte lineages based on their module scores. This lineage assignment was used for all subsequent lineage-stratified analyses, including the RNA Features approach described below.
2. <u>RNA Features approach</u>: Homotypic doublets were removed as upper tail outliers in the distribution of the number of transcripts (nCount_RNA) and genes (nFeature_RNA) per droplet. Because baseline RNA content is different between lineages and experimental runs, the filter was defined as a global top quantile rather than an absolute cutoff. For a given quantile, the corresponding cutoffs were computed separately for each sequencing run (run1 and run2) and lineage (T cell and monocyte), and droplets exceeding the threshold for either nCount_RNA or nFeature_RNA were removed. To select this global quantile, we used a grid search of top-expression quantiles ranging from 0% to 5% (step size 0.05%). Because the in-contact population had the highest frequency of inferred intra-sample doublets, the quantile was chosen to match the number of droplets removed from this population. For each candidate quantile in the grid search, the number of outlier droplets removed from the in-contact population was summed across lineages within each run and compared to the target number, and the total absolute discrepancy across both runs was calculated. Minimizing this discrepancy yielded an optimal threshold at 4.1-4.2%, and 4.1% was selected as the global cutoff (**Figure S2D**); droplets above this threshold in nCount_RNA or nFeature_RNA were removed (**Figure 2D**).

#### Low-quality cells removal

Low-quality droplets were removed using a symmetric lower threshold. Within each lineage and run, cells below the bottom 1% quantile of nCount_RNA or nFeature_RNA were discarded (**Figures S2H, S2I**).

#### Dataset split into T cells and monocytes

After removal of multi-aggregates, inter-sample doublets, intra-sample doublets, and low-quality droplets, the retained cells were re-clustered to generate the final embedding used for all downstream analyses. Normalization and variance stabilization were again performed using the function ‘SCTransform’ (v2 workflow, regressing out percent.mt), followed by running the function ‘FindNeighbors’ (setting dims = 1:30, k.param = 50) and the function ‘FindClusters’ (setting resolution = 0.4) on the SCT assay. Cells were embedded in two dimensions using the function ‘RunUMAP’ (setting dims = 1:30, metric = “euclidean”, seed.use = 16). T cell and monocyte clusters were annotated based on the T cell and monocyte module scores and split from the original Seurat Object into one T cell and one monocyte Seurat object by extracting raw RNA counts.

For dimensionality reduction, clustering, and visualization steps, the same functions and parameters were used except that the resolution was set to 0.6 and 0.3 for the T cell and the monocyte Seurat objects, respectively. To determine the resolution for identifying clusters, R package clustree (v0.5.1) (28) was used. NormalizeData and ScaleData functions were used to generate data slot and scale.data slot, respectively, from raw RNA counts for both subset Seurat objects. Top genes for each cluster were extracted using the FindAllMarkers function on SCT assay and data slot (setting min.pct = 0.25, logfc.threshold = 0.25, test.use = “wilcox”), selecting only positive genes (adjusted p-value < 0.05).

#### Pseudobulk differential expression analysis

Pseudobulk differential expression (DE) analysis was performed on raw RNA counts. To control for total cell number imbalances across participants, cells were first downsampled within each participant to equalize cell counts between the two compared conditions. Then, raw RNA counts were aggregated across individual participant-condition pairs to generate pseudobulk count matrices. Genes were filtered to retain those expressed in at least 10% of cells in either group and with a total cumulative pseudobulk count of at least 10 reads across all samples. DE analysis was performed using R package DESeq2 (v 1.42.1) (29). To account for participant-to-participant variability, a paired design formula (∼ ParticipantID + Gate_Doublet_collapsed) was implemented using the function ‘DESeqDataSetFromMatrix’ and the function ‘DESeq’. Results were extracted using the function ‘results’ (setting independentFiltering = FALSE). DE genes were defined as those with an Benjamini-Hochberg (BH) adjusted p-value < 0.05 and an absolute value of log2 fold-change (FC) > 0.2.

#### Gene set enrichment analysis (GSEA)/Gene ontology (GO) analysis

To identify biological pathways enriched between conditions, gene set enrichment analysis (GSEA) was performed using R package clusterProfiler (v4.10.1) (30). All genes were ranked in descending order based on log2 FC values. GSEA was conducted against Gene Ontology (GO) Biological Process (BP) terms using the function ‘gseGO’ (setting OrgDb = org.Hs.eg.db, keyType = “SYMBOL”, ont = “BP”, minGSSize = 25, maxGSSize = 500, and pvalueCutoff = 0.05). To remove redundant GO terms, the resulting pathway enrichments were simplified using the function ‘simplify’ (setting cutoff = 0.5, by = “p.adjust”, and select_fun = min). GO terms with an adjusted p-value < 0.05 were considered significantly enriched. GO BP enrichment analysis was performed using enrichR (https://maayanlab.cloud/Enrichr/) (31) for BP terms. Redundant terms with overlapping gene sets were removed. Pathways with a BH-adjusted p-value < 0.05 were considered significantly enriched.

#### Gene module score calculation

Gene module scores were calculated using the R package UCell (v2.6.2) (32). For defined gene sets, module scores were generated using the function ‘AddModuleScore_UCell’ (setting maxRank = 3000, and chunk.size = 5000). This rank-based scoring method calculates single-cell signature scores based on Mann-Whitney U statistics relative to the top 3,000 expressed genes within each cell.

#### T cell-monocyte intermixing model

Within each participant, cells were randomly downsampled to equalize cell numbers across singlet T-cell (*T_singlet_*), in-contact T-cell (*T_in-contact_*), and singlet monocyte (*M_singlet_*) populations. Pseudobulk count matrices were then constructed by summing raw RNA counts across downsampled cells per participant for each population. Simulated in-contact T-cell profiles (*T_sim_*) were modeled across a fine grid of monocyte contamination fractions *x (0 ≤ x ≤ 1)* using the formula: *T_sim_* (*x*) = *T_singlet_* + *x* * *M_singlet_*. For each candidate fraction *x*, paired DE analysis between *T_sim_* (*x*) and *T_in-contact_* and T_in-contact_ was conducted across participants using R package DESeq2. Size factors and dispersions were calculated using the function ‘estimateSizeFactors’ and the function ‘estimateDispersionsGeneEst’, followed by testing using the function ‘nbinomWaldTest’ with a paired design formula (∼ participant + condition). DE genes between *T_sim_* (*x*) and *T_in-contact_* cells were identified using the function results (setting independentFiltering = FALSE) with thresholds of BH-adjusted padj < 0.05 and |log2FoldChange| > 0.2. The optimal monocyte contamination fraction 0.41% was defined as that minimized the total number of DE genes between *T_sim_* (*x*) and *T_in-contact_* (**Figure S3A**).

#### DICE gene expression

Gene lists were queried against the DICE (Database of Immune Cell Expression) (18) CellTypeScore web tool. Gene lists were submitted to extract baseline expression levels in Transcripts Per Million (TPM) across reference immune cell types. To allow direct comparison of cell-type distribution across genes, min-max normalization was applied to rescale TPM values between 0 and 1 for each gene:

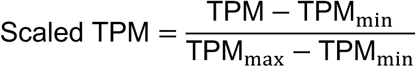

where TPM_min_ and TPM_max_ represent the minimum and maximum expression levels observed for a specific gene across all cell types.

#### Coincidental curve prediction

For singlet, coincidental, and in-contact populations, UCell scores were binned into 60 equal-width intervals spanning the score range observed across combined populations, and density histograms normalized to unit area were computed ( *d_S_*, *d_I_*, *d_C_* for singlet, in-contact, and coincidental, respectively). The coincidental density was modeled as a mixture of singlet and in-contact densities, *d_C_* = *α* · *d_S_* + (1 − *α*) · *d_I_*, and the mixing coefficient α was estimated by ordinary least-squares (*d_C_* − *d_I_*) = *α* · (*d_S_* − *d_I_*), with α constrained to the interval [0,1]. Goodness of fit for the coincidental population was quantified as the coefficient of determination (*R*²) between the observed density and the predicted mixture density *α* · *d_S_* + (1 − *α*) · *d_I_*.

#### Dual lineage droplet analysis

For dual-lineage droplet analysis, two droplet types were isolated and combined: double-positive (DP) T cell-monocyte doublets identified during the intra-sample doublet removal step with the Dual Expression approach (**Figure 2C**), and the T cell-like monocyte cluster (c7) from the monocyte dataset (**Figures 5A, S5B**). Raw counts from each droplet were retained with unaggregated genes and were subsequently normalized and variance-stabilized using the function ‘NormalizeData’ and the function ‘SCTransform’ (regressing out percent.mt) to preserve full gene-level resolution.

### Single-cell TCR sequencing data analysis

#### Clonal expansion analysis

TCR data was analyzed using R package scRepertoire (v2.5.2) (33). We first constructed a TCR genes table combining all Cell Ranger output files filtered_contig_annotations.csv for each sample using the function ‘combineTCR’ (setting removeNA = TRUE, removeMulti = FALSE, filterMulti = FALSE, filterNonproductive = TRUE). Next, clusters corresponding to γδ T cells and CD4^-^CD8^-^ T cells were excluded from downstream TCR analyses because their TCRαβ detection rates fell below 50% (**Figure S4A**). In the remaining clusters, we filtered cells that had both TCRα and TCRβ sequences detected, resulting in 14,796 cells across all 20 participants, combining singlet and in-contact T cells (**Table S5** for TCRαβ sequences retrieved for each cell).

For each participant, cells were randomly downsampled per singlet and in-contact categories using the function ‘slice_sample’ from R package dplyr (v1.1.4) to match the smallest cell count available for that participant across categories. Within each bootstrap iteration, clonotypes were defined based on unique TCR amino acid sequences, and clonotypes with two or more cells were classified as expanded. The percentage of expanded clonotypes was calculated as the number of expanded clonotypes divided by the total number of unique clonotypes per participant and category and averaged across all 100 bootstrap iterations. In parallel, within each bootstrap iteration, diversity indices, including Shannon index, Chao1 estimator, Gini-Simpson index, and normalized entropy, were calculated using the function ‘clonalDiversity’ (setting cloneCall = “aa”, chain = “both”, and exportTable = TRUE) from R package scRepertoire. Metric values were averaged across the 20 bootstrap iterations per participant and category.

#### Gene usage analysis

TCR V-gene usage was analyzed from paired α/β chain contigs. For each cell, the TRAV and TRBV genes were extracted from the α and β chain annotations, and allelic subgroups were aggregated to the gene level. Within each T cell meta-cluster, the relative usage of each gene for singlet and in-contact T cells was calculated by the number of cells using that gene divided by the total number of cells with both α and β chain sequenced in that group.

#### Antigen-specific analysis

Clusters corresponding to γδ T cells and CD4^-^CD8^-^ T cells were excluded from antigen-specific analyses. In the remaining clusters, we kept cells that had at least one TCRβ sequence detected, resulting in 15,995 cells across all 20 participants, combining singlet and in-contact T cells (**Table S6**). The antigen specificity of retrieved TCR CDR3β sequences was evaluated by matching them against three reference databases:

1. TCRMatch (v1.3) (15) to identify known epitope specificities within the Immune Epitope Database (IEDB, https://www.iedb.org, setting match score threshold = 0.97) (34);
2. A collated database of 32,869 unique Mtb-reactive CDR3β sequences compiled by Turner et al. (14) (encompassing 422 sequences from VDJdb, 16,249 sequences from McPAS-TCR, and 16,475 sequences from a previous study (35));
3. A panel of 1,392 candidate Mtb-reactive CDR3β metaclones derived using the Metaclonotypist computational pipeline developed by Turner et al. (14).

For databases 2 and 3, pairwise sequence distances between unique query CDR3β sequences and reference Mtb sequences were computed using the pwseqdist Python package (v0.6, the function pw.metrics.nb_vector_tcrdist via pw.apply_pairwise_rect). A CDR3β sequence was defined as Mtb-reactive if it has a TCRdist score of 4 or less to any reference Mtb sequence.

For each participant, cell counts per singlet and in-contact categories were downsampled to match the minimum cell count available across both categories for that specific participant. Within each bootstrap iteration, the percentage of Mtb-reactive unique clonotypes (unique Mtb-reactive CDR3β sequences divided by total unique CDR3β sequences) and the percentage of Mtb-reactive cells were calculated for each participant and gate. Metric values were averaged across all 100 bootstrap iterations per participant and gate for downstream paired statistical testing across participants.

### Quantification and Statistical analysis

Statistical analyses were performed using ggsignif (v0.6.4) from R. Paired datasets were compared using non-parametric Wilcoxon tests. P values less than 0.05 were considered significant and 2-tailed analyses were performed. Correction for multiple comparisons was performed with Benjamini-Hochberg (BH) correction when applicable. For all the boxplots, the lower, median, and upper edges represent the 25th, 50th, and 75th percentile; the length of the upper and lower whiskers is 1.5 times the interquartile range. To evaluate differences in the proportion of participants with detectable antigen-specific TCRs (>0%) between paired singlet and in-contact cells across matched participants (n=20), McNemar’s chi-square test was performed on the 2 x 2 contingency table of paired status with df = 1 using function ‘mcnemar.test’ (setting alternative = “greater”) from R package stats.

## Data availability

Single-cell RNA and TCR sequencing data are available in GEO and Immport under accession number [Pending].

## Author contributions

Conceptualization (BP, JGB); Methodology (CK, JGB); Investigation (NK, CK, JGB); Formal analysis (NK, JGB); Resources (TJS); Writing – original draft (NK, JGB); Writing – Review and editing (NK, CK, TJS, BP, JGB); Funding acquisition (BP, JGB).

## Acknowledgments

We thank the Flow cytometry, Sequencing, and Bioinformatics core facilities at La Jolla Institute for Immunology for technical assistance with cell sorting, sequencing, and data analysis. We thank Drs. Gisele Baracho, Aaron Tyznik and Aaron Middlebrook (Waters Biosciences) for insightful discussions regarding image-guided gating of T cell-monocyte doublets. We also express our gratitude to Drs. Robert H. Gilman and Mayuko Saita for providing participant PBMC samples from the Universidad Peruana Cayetano Heredia (Peru).

## References

1. Kang N, Kim C, Peters B, Burel JG. Decoding Immune Cell Doublets in Humans: From Artifacts to Insights. Cytom Part J Int Soc Anal Cytol. 2026 Aug 14. doi:10.1002/cyto.a.70060 PubMed PMID: 42601345.

2. Burel JG, Pomaznoy M, Lindestam Arlehamn CS, Weiskopf D, da Silva Antunes R, Jung Y, et al. Circulating T cell-monocyte complexes are markers of immune perturbations. eLife. 2019 Jun 25;8:e46045. doi:10.7554/eLife.46045 PubMed PMID: 31237234; PubMed Central PMCID: PMC6592685.

3. Gil-Manso S, Miguens Blanco I, López-Esteban R, Carbonell D, López-Fernández LA, West L, et al. Comprehensive Flow Cytometry Profiling of the Immune System in COVID-19 Convalescent Individuals. Front Immunol. 2021;12:793142. doi:10.3389/fimmu.2021.793142 PubMed PMID: 35069575; PubMed Central PMCID: PMC8771913.

4. Lin J, Bai S, He L, Yang Y, Li X, Luo L, et al. Cytotoxic Lymphocyte-Monocyte Complex Reflects the Dynamics of Coronavirus Disease 2019 Systemic Immune Response. J Infect Dis. 2024 Jul 25;230(1):5–14. doi:10.1093/infdis/jiae048 PubMed PMID: 39052699.

5. Cortese M, Hagan T, Rouphael N, Wu SY, Xie X, Kazmin D, et al. System vaccinology analysis of predictors and mechanisms of antibody response durability to multiple vaccines in humans. Nat Immunol. 2025 Jan;26(1):116–30. doi:10.1038/s41590-024-02036-z PubMed PMID: 39747435.

6. Obare LM, Simmons J, Oakes J, Zhang X, Nochowicz C, Priest S, et al. CD3+ T-cell: CD14+ monocyte complexes are dynamic and increased with HIV and glucose intolerance. J Immunol Baltim Md 1950. 2025 Feb 28;214(3):516–31. doi:10.1093/jimmun/vkae054 PubMed PMID: 40073149; PubMed Central PMCID: PMC11952877.

7. Polasky C, Steffen A, Loyal K, Lange C, Bruchhage KL, Pries R. Redistribution of Monocyte Subsets in Obstructive Sleep Apnea Syndrome Patients Leads to an Imbalanced PD-1/PD-L1 Cross-Talk with CD4/CD8 T Cells. J Immunol Baltim Md 1950. 2021 Jan 1;206(1):51–8. doi:10.4049/jimmunol.2001047 PubMed PMID: 33268482.

8. Kang N, Chawla A, Hillman H, Tippalagama R, Kim C, Mikulski Z, et al. T cell monocyte complexes exhibit distinct immune signatures during infection. iScience. 2025 Sep 19;28(9):113432. doi:10.1016/j.isci.2025.113432 PubMed PMID: 40989028; PubMed Central PMCID: PMC12450731.

9. Burel JG, Pomaznoy M, Lindestam Arlehamn CS, Seumois G, Vijayanand P, Sette A, et al. The Challenge of Distinguishing Cell-Cell Complexes from Singlet Cells in Non-Imaging Flow Cytometry and Single-Cell Sorting. Cytom Part J Int Soc Anal Cytol. 2020 Nov;97(11):1127–35. doi:10.1002/cyto.a.24027 PubMed PMID: 32400942; PubMed Central PMCID: PMC7666012.

10. Schraivogel D, Kuhn TM, Rauscher B, Rodríguez-Martínez M, Paulsen M, Owsley K, et al. High-speed fluorescence image-enabled cell sorting. Science. 2022 Jan 21;375(6578):315–20. doi:10.1126/science.abj3013 PubMed PMID: 35050652; PubMed Central PMCID: PMC7613231.

11. Kuhn TM, Paulsen M, Cuylen-Haering S. Accessible high-speed image-activated cell sorting. Trends Cell Biol. 2024 Aug;34(8):657–70. doi:10.1016/j.tcb.2024.04.007 PubMed PMID: 38789300.

12. Giladi A, Cohen M, Medaglia C, Baran Y, Li B, Zada M, et al. Dissecting cellular crosstalk by sequencing physically interacting cells. Nat Biotechnol. 2020 May;38(5):629–37. doi:10.1038/s41587-020-0442-2 PubMed PMID: 32152598.

13. Kim C, Hinz D, Middlebrook A, Tyznik AJ, D’Cruz LM, Scriba T, et al. A high-resolution blood immune cell doublet atlas via imaging spectral cytometry. bioRxiv; 2026 [cited 2026 Sep 18]. Available from: http://biorxiv.org/lookup/doi/10.64898/2026.09.03.749150 doi:10.64898/2026.09.03.749150

14. Turner CT, Tiffeau-Mayer A, Rosenheim J, Chandran A, Saxena R, Zhang P, et al. Evolution of the tuberculin skin test reveals generalisable Mtb-reactive T cell metaclones. Nat Commun. 2026 Jan 21;17(1):1900. doi:10.1038/s41467-026-68678-9 PubMed PMID: 41565657; PubMed Central PMCID: PMC12923891.

15. Chronister WD, Crinklaw A, Mahajan S, Vita R, Koşaloğlu-Yalçın Z, Yan Z, et al. TCRMatch: Predicting T-Cell Receptor Specificity Based on Sequence Similarity to Previously Characterized Receptors. Front Immunol. 2021;12:640725. doi:10.3389/fimmu.2021.640725 PubMed PMID: 33777034; PubMed Central PMCID: PMC7991084.

16. Nelson CE, Kauffman KD, Osum KC, Sakai S, Buchanan J, Chanchu JM, et al. Mycobacterium tuberculosis peptide-specific T cells in pulmonary granulomas display broad effector functions. iScience. 2025 Dec 19;28(12):114034. doi:10.1016/j.isci.2025.114034 PubMed PMID: 41438074; PubMed Central PMCID: PMC12721174.

17. Tippalagama R, Trevizani R, Chihab LY, Chawla A, Fung K, Greenbaum J, et al. Peptide-driven identification of TCRs reveals dynamics and phenotypes of CD4 T cells in tuberculosis. J Immunol. 2026 Feb 9;215(2):vkaf287. doi:10.1093/jimmun/vkaf287 PubMed PMID: 41241821; PubMed Central PMCID: PMC12718515.

18. Schmiedel BJ, Singh D, Madrigal A, Valdovino-Gonzalez AG, White BM, Zapardiel-Gonzalo J, et al. Impact of Genetic Polymorphisms on Human Immune Cell Gene Expression. Cell. 2018 Nov 29;175(6):1701–1715.e16. doi:10.1016/j.cell.2018.10.022 PubMed PMID: 30449622; PubMed Central PMCID: PMC6289654.

19. Céspedes PF, Jainarayanan A, Fernández-Messina L, Valvo S, Saliba DG, Kurz E, et al. T-cell trans-synaptic vesicles are distinct and carry greater effector content than constitutive extracellular vesicles. Nat Commun. 2022 Jun 16;13(1):3460. doi:10.1038/s41467-022-31160-3 PubMed PMID: 35710644; PubMed Central PMCID: PMC9203538.

20. Mittelbrunn M, Gutiérrez-Vázquez C, Villarroya-Beltri C, González S, Sánchez-Cabo F, González MÁ, et al. Unidirectional transfer of microRNA-loaded exosomes from T cells to antigen-presenting cells. Nat Commun. 2011;2:282. doi:10.1038/ncomms1285 PubMed PMID: 21505438; PubMed Central PMCID: PMC3104548.

21. Haimovich G, Ecker CM, Dunagin MC, Eggan E, Raj A, Gerst JE, et al. Intercellular mRNA trafficking via membrane nanotube-like extensions in mammalian cells. Proc Natl Acad Sci U S A. 2017 Nov 14;114(46):E9873–82. doi:10.1073/pnas.1706365114 PubMed PMID: 29078295; PubMed Central PMCID: PMC5699038.

22. Escoll P, Buchrieser C. Metabolic reprogramming of host cells upon bacterial infection: Why shift to a Warburg-like metabolism? FEBS J. 2018 Jun;285(12):2146–60. doi:10.1111/febs.14446 PubMed PMID: 29603622.

23. Hillman H, Khan N, Singhania A, Dubelko P, Soldevila F, Tippalagama R, et al. Single-cell profiling reveals distinct subsets of CD14+ monocytes drive blood immune signatures of active tuberculosis. Front Immunol. 2022;13:1087010. doi:10.3389/fimmu.2022.1087010 PubMed PMID: 36713384; PubMed Central PMCID: PMC9874319.

24. Mortensen R, Arlehamn CSL, Coler RN, Gerner MY, Goletti D, Lewinsohn DA, et al. T cell-macrophage interactions in tuberculosis: What we’ve got here is failure to communicate. J Intern Med. 2026 Jan;299(1):44–65. doi:10.1111/joim.70028 PubMed PMID: 41273222; PubMed Central PMCID: PMC13289048.

25. Habtamu M, Abrahamsen G, Aseffa A, Andargie E, Ayalew S, Abebe M, et al. High-throughput analysis of T cell-monocyte interaction in human tuberculosis. Clin Exp Immunol. 2020 Aug;201(2):187–99. doi:10.1111/cei.13447 PubMed PMID: 32348546; PubMed Central PMCID: PMC7366737.

26. Zheng GXY, Terry JM, Belgrader P, Ryvkin P, Bent ZW, Wilson R, et al. Massively parallel digital transcriptional profiling of single cells. Nat Commun. 2017 Jan 16;8:14049. doi:10.1038/ncomms14049 PubMed PMID: 28091601; PubMed Central PMCID: PMC5241818.

27. Hao Y, Stuart T, Kowalski MH, Choudhary S, Hoffman P, Hartman A, et al. Dictionary learning for integrative, multimodal and scalable single-cell analysis. Nat Biotechnol. 2024 Feb;42(2):293–304. doi:10.1038/s41587-023-01767-y PubMed PMID: 37231261; PubMed Central PMCID: PMC10928517.

28. Zappia L, Oshlack A. Clustering trees: a visualization for evaluating clusterings at multiple resolutions. GigaScience. 2018 Jul 1;7(7):giy083. doi:10.1093/gigascience/giy083 PubMed PMID: 30010766; PubMed Central PMCID: PMC6057528.

29. Love MI, Huber W, Anders S. Moderated estimation of fold change and dispersion for RNA-seq data with DESeq2. Genome Biol. 2014;15(12):550. doi:10.1186/s13059-014-0550-8 PubMed PMID: 25516281; PubMed Central PMCID: PMC4302049.

30. Yu G, Wang LG, Han Y, He QY. clusterProfiler: an R package for comparing biological themes among gene clusters. Omics J Integr Biol. 2012 May;16(5):284–7. doi:10.1089/omi.2011.0118 PubMed PMID: 22455463; PubMed Central PMCID: PMC3339379.

31. Kuleshov MV, Jones MR, Rouillard AD, Fernandez NF, Duan Q, Wang Z, et al. Enrichr: a comprehensive gene set enrichment analysis web server 2016 update. Nucleic Acids Res. 2016 Jul 8;44(W1):W90–97. doi:10.1093/nar/gkw377 PubMed PMID: 27141961; PubMed Central PMCID: PMC4987924.

32. Andreatta M, Carmona SJ. UCell: Robust and scalable single-cell gene signature scoring. Comput Struct Biotechnol J. 2021;19:3796–8. doi:10.1016/j.csbj.2021.06.043 PubMed PMID: 34285779; PubMed Central PMCID: PMC8271111.

33. Borcherding N, Bormann NL, Kraus G. scRepertoire: An R-based toolkit for single-cell immune receptor analysis. F1000Research. 2020;9:47. doi:10.12688/f1000research.22139.2 PubMed PMID: 32789006; PubMed Central PMCID: PMC7400693.

34. Vita R, Blazeska N, Marrama D, IEDB Curation Team Members, Duesing S, Bennett J, et al. The Immune Epitope Database (IEDB): 2024 update. Nucleic Acids Res. 2025 Jan 6;53(D1):D436–43. doi:10.1093/nar/gkae1092 PubMed PMID: 39558162; PubMed Central PMCID: PMC11701597.

35. Musvosvi M, Huang H, Wang C, Xia Q, Rozot V, Krishnan A, et al. T cell receptor repertoires associated with control and disease progression following Mycobacterium tuberculosis infection. Nat Med. 2023 Jan;29(1):258–69. doi:10.1038/s41591-022-02110-9 PubMed PMID: 36604540; PubMed Central PMCID: PMC9873565.

