## Supplemental material for "Single-cell profiling of physically interacting immune cells using image-guided sorting"

Content

*Figures S1 to S7*

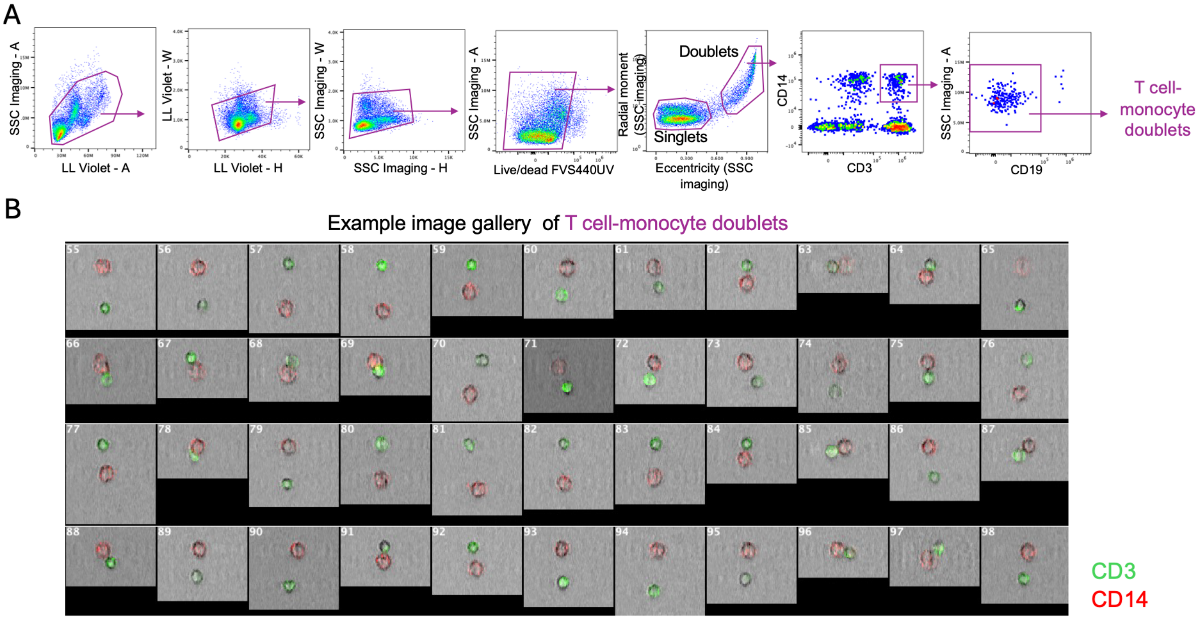


**Figure S1: Imaging cell sorting reveals heterogeneity within blood-derived T cell-monocyte doublets.** (A) Gating strategy to identify T cell-monocyte doublets using the imaging cell sorter BD FACS Discover™ S8 in PBMC. (B) Example image gallery of T cell-monocyte doublets as gated in panel A.

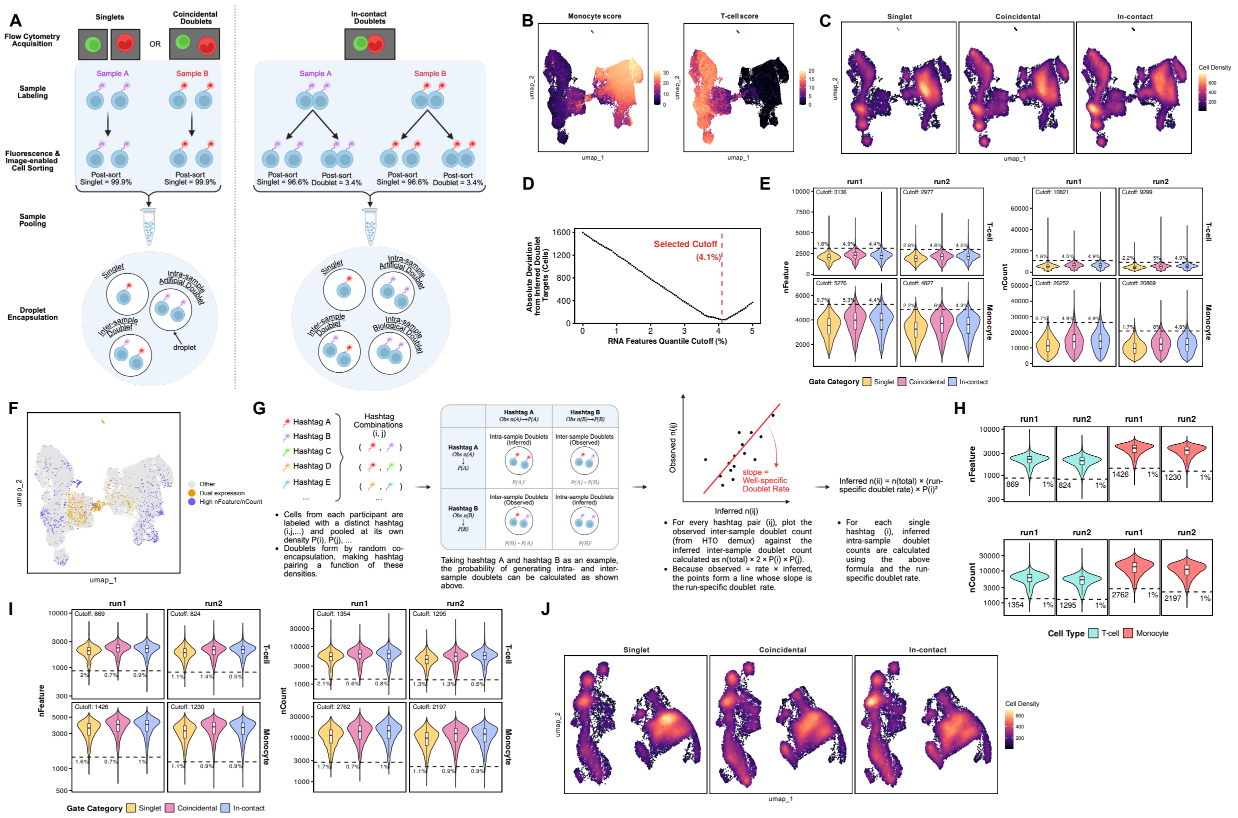

**Figure S2: Supplementary analyses for the computational identification and removal of inter- and intra-sample doublets.** (A) Schematic showing the different doublet types expected to be generated from three image-gated categories of T cell-monocyte pairs after sorting and single-cell droplet-based sequencing. Pre-QC UMAP plots showing (B) T cell and monocyte module scores and (C) cell density across image-gated categories. (D) Line plot displaying optimization of the upper RNA feature quantile cutoff (0-5%, step size 0.05%) based on absolute deviation from target inferred doublet counts. The red dashed line marks the optimal cutoff (4.1%). (E) Violin plots showing number of genes (nFeature, left) and transcripts (nCount, right) distributions across sequencing runs (run 1, run 2), cell lineages (T-cell, monocyte), and image-gated sorted categories (singlet, coincidental, in-contact), with upper cutoff thresholds (top 4.1% quantile) indicated. (F) Pre-QC UMAP highlighting doublets identified by dual expression (gold) or RNA features (blue) approaches. (G) Schematic outlining the inference of artificial intra-sample doublet frequencies from inter-sample doublet (HTO demux) co-encapsulation rates. Violin plots showing nFeature (top) and nCount (bottom) distributions across sequencing runs and cell lineages, presented (H) overall or (I) split by image-gated categories, with lower cutoff thresholds (bottom 1% quantile) indicated. (J) Post-QC UMAP plot displaying cell density across image-gated categories. For E, H, and I: Black dashed lines denote quantile cutoffs with exact numeric values annotated; percentage labels indicate removed outliers per category; inner white boxplots represent the median and IQR. For J, data derived from 69,640 post-QC droplets; for B, C, and F, data derived from 76,918 pre-QC droplets; for all, data derived from n=20 PBMC samples.

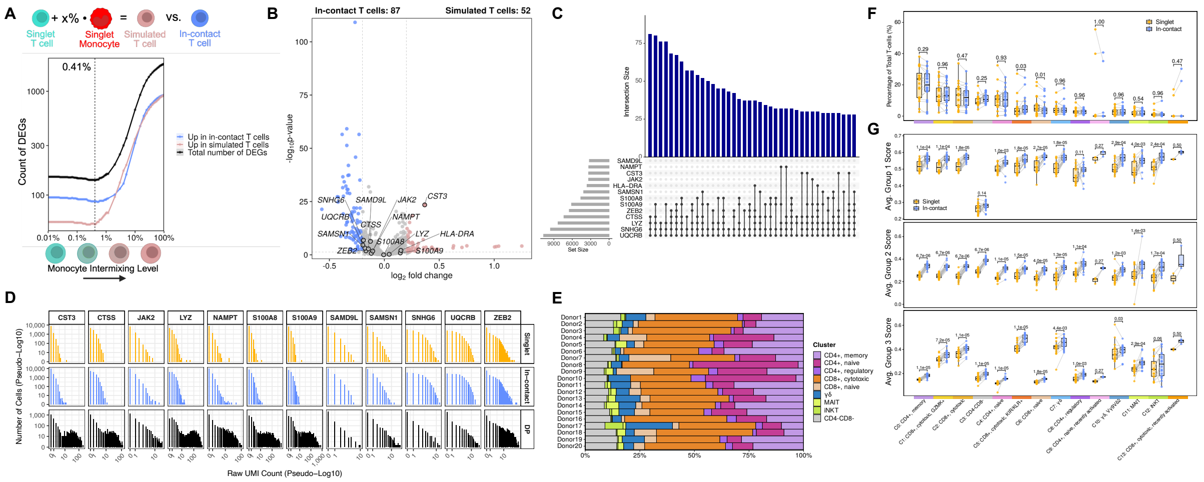


**Figure S3: Supplementary transcriptomic analyses of in-contact T cells.** (A) T cell-monocyte intermixing model. Plotted lines represent genes enriched in in-contact T cells (blue), genes enriched in simulated T cells (coral), and the total number of DEGs (black) when comparing simulated T cells with a gradient of intermixing monocyte contamination percentages vs in-contact T cells. The vertical black dashed line marks the monocyte contamination fraction that most resemble in-contact T cells. (B) Volcano plot of DEGs comparing simulated T cells at 0.41% monocyte intermixing level (T-$M_{0.41\%}$) versus in-contact T cells. Genes with BH-adjusted $p < 0.05$ and |average log2 FC| > 0.2 are colored by the category in which they are upregulated (simulated T cells, pink; in-contact T cells (blue). Genes significantly upregulated in in-contact T cells compared to singlet T cells (T.90 signature, Figure 3A) are labeled with their gene name. (C) UpSet plot showing single-cell co-expression combinations for 13 monocyte-associated genes in in-contact T cells. The top bar plot indicates the number of single cells co-expressing each specific gene combination (ordered by frequency), while the left bar plot shows total cell counts expressing each individual gene. Connected dots in the matrix mark the specific genes included in each co-expression set. (D) Histograms with the per-cell raw UMI counts for 13 monocyte-associated genes in singlet (gold) T cells, in-contact T cells (blue) and DP T cell-monocyte droplets (black). Both axes are shown in a pseudo-log10 scale. (E) Stacked bar plot showing the frequency of T cell meta-clusters per participant (normalized to 100%). (F) Individual cluster composition differences between singlet and in-contact T cells. (G) Boxplots of per-participant average UCell module scores for three gene groups derived from the T.90 in-contact signature across T cell clusters. For both F and G, paired per-participant points are shown for singlet (gold) and in-contact (blue) T cells, with grey lines connecting matched participants; boxplots show the median and IQR. Gates were compared within each subset by paired two-sided Wilcoxon signed-rank test with BH adjustment (adjusted p-values above brackets). Data were derived from 12,487 in-contact, 9,272 singlet T cells and 2,455 DP droplets, from n=20 PBMC samples of patients with ATB collected at diagnosis.

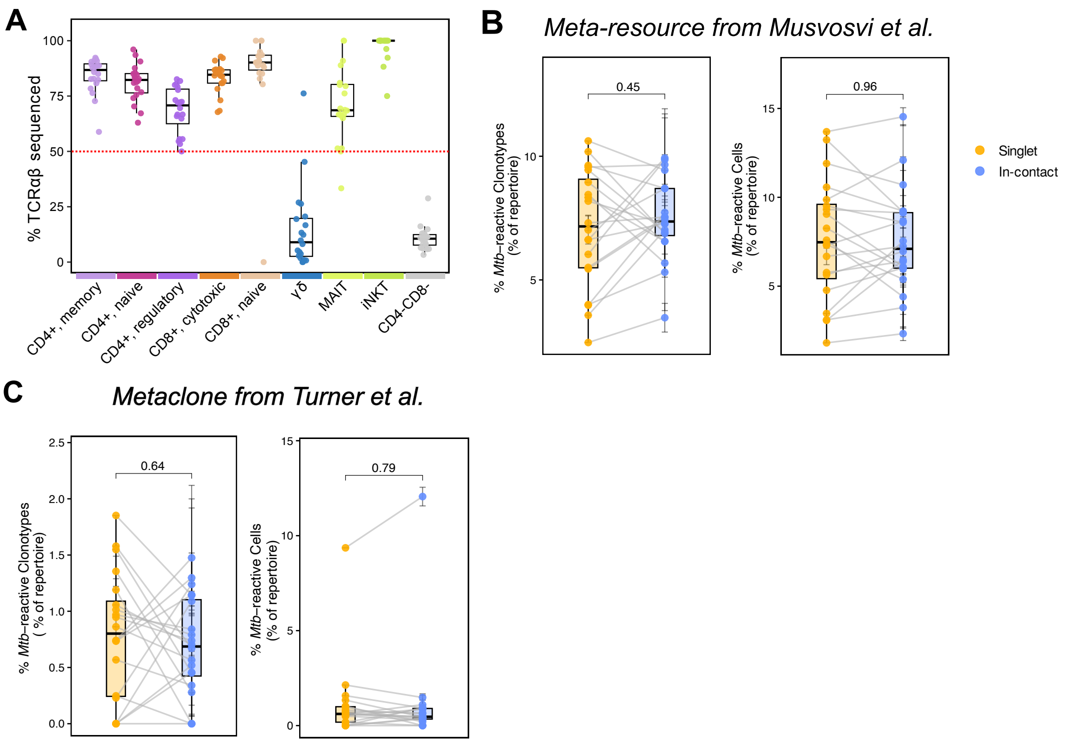


**Figure S4: Supplementary TCR repertoire analyses of in-contact T cells.** (A) Dotplot showing the percentage of cells with TCRαβ across T cell meta-clusters. Dots represent individual participants, and boxplots display the participant-level median and IQR across combined singlet and in-contact T cells. (B) Proportions of Mtb-reactive TCR clonotypes relative to total TCRβ clonotypes (left) and Mtb-reactive cells relative to total cells with TCRβ (right), identified using the collated database from Turner et al., 2026 (see method for details). (C) Proportions of Mtb-reactive TCR clonotypes relative to total TCRβ clonotypes (left) and Mtb-reactive cells relative to total cells with TCRβ (right), identified using the Mtb metaclones from Turner et al., 2026 (see method for details). For B and C, values represent participant-level mean percentages derived from 100 bootstrap downsampling iterations. Boxplots display the median and IQR, and gray lines connect matched participant samples. Statistical significance was evaluated using paired one-sided Wilcoxon signed-rank tests; p-values are indicated above brackets. For all panels, data are derived from 15,995 T cells with TCRβ sequences detected (excluding γδ and $CD4^{-}CD8^{-}$ metaclusters) from n=20 PBMC samples of patients with ATB collected at diagnosis.

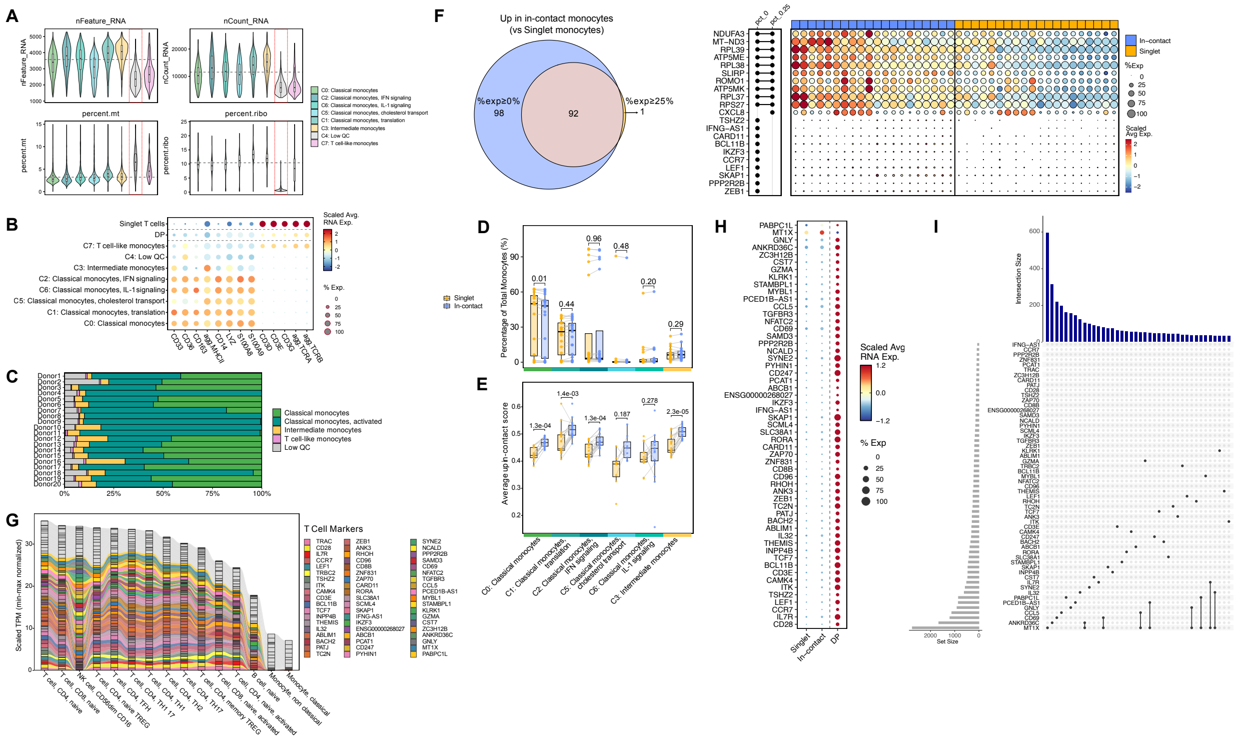

**Figure S5: Supplementary transcriptomic analyses of in-contact monocytes.** (A) Violin plots showing number of genes (nFeature_RNA, upper left), transcripts (nCount_RNA, upper right), percentage of mitochondria genes (percent.mt, lower left) and ribosomal genes (percent.ribo, lower right) distributions across monocyte clusters. Horizontal dashed lines show the average value across all clusters. (B) Dot plot showing canonical T cell genes and monocyte genes across singlet T cells, DP T cell-monocyte droplets, and monocyte clusters. (C) Stacked bar plot showing the frequencies of monocyte clusters per participant (normalized to 100%). (D) Monocyte cluster composition differences between in-contact (blue) and singlet (gold) monocytes paired by sample. (E) Boxplots of per-participant average UCell module scores for 93 DEGs significantly enriched in in-contact monocytes across monocyte clusters. For D and E, paired per-participant points are shown for singlet (gold) and in-contact (blue) monocytes, with grey lines connecting matched participants; boxplots show the median and IQR. Gates were compared within each subset by paired two-sided Wilcoxon signed-rank test with BH adjustment (adjusted p-values above brackets). (F) Comparison of DEGs upregulated in in-contact (blue) versus singlet (gold)

monocytes under two percent-expression cutoffs (0% and 25%). (Left) Venn diagram showing the number of upregulated DEGs shared or unique between the two cutoffs (Right) Dot plot of the top upregulated DEGs across individual participants, split by in-contact (blue) and singlet (gold) monocytes. Dot color intensity represents scaled average expression and dot size the percentage of expressing cells within each participant/gate group. (G) Alluvial plot showing min-max normalized TPM (y-axis) from the DICE database of the 98 upregulated DEGs uniquely upregulated in in-contact monocytes at a 0% expression fraction cutoff. Gene expression values are scaled between 0 and 1. Cell types on the x-axis are ordered by cumulative scaled expression scores. Colored bars represent genes with a log2 ratio of mean T cell to monocyte expression > 2. (H) Dot plot showing T cell-associated upregulated DEGs across singlet monocytes, in-contact monocytes and DP T cell-monocyte droplets. (I) UpSet plot showing single-cell co-expression combinations for T cell-associated upregulated DEGs in in-contact monocytes. The top bar plot indicates the number of single cells co-expressing each specific gene combination (ordered by frequency), while the left bar plot shows total cell counts expressing each individual gene. Connected dots in the matrix mark the specific genes included in each co-expression set. Data were derived from 11,915 in-contact, 10,448 singlet monocytes and 2,455 DP droplets, from n=20 PBMC samples of patients with ATB collected at diagnosis.

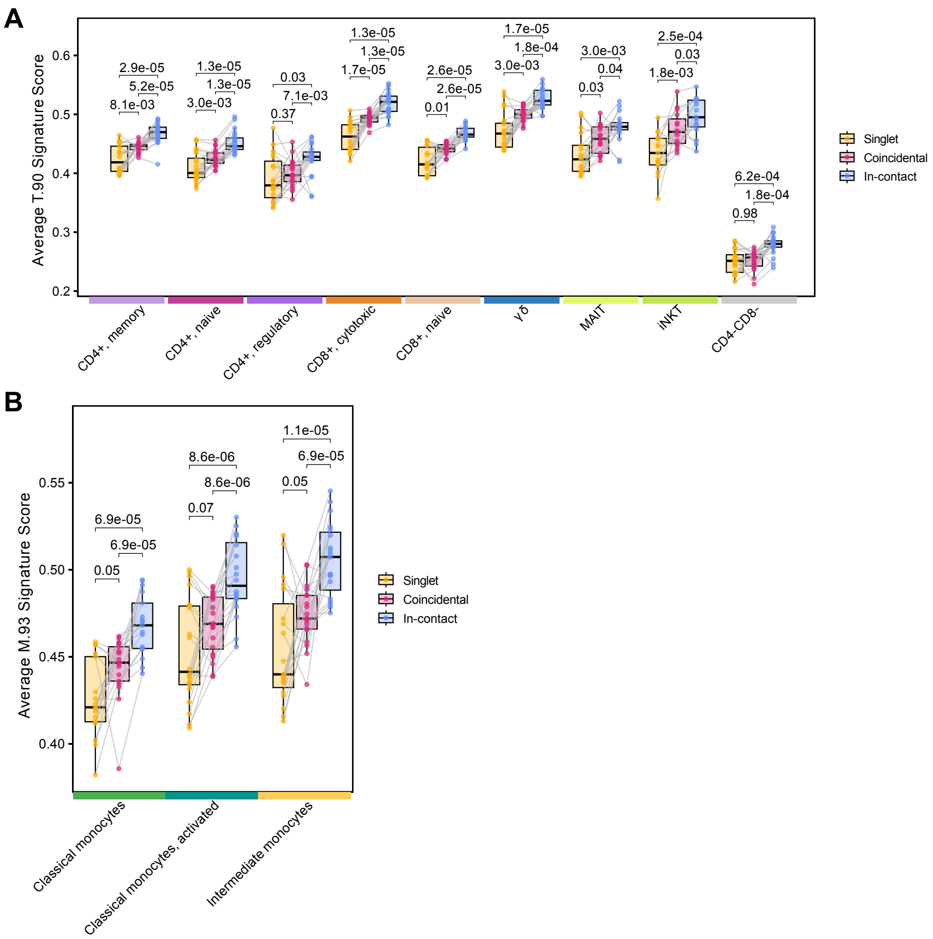


**Figure S6: Supplementary transcriptomic analyses of coincidental T cells and monocytes.** (A) Boxplots of per-participant average UCell module scores for (A) 90 DEGs upregulated in in-contact T cells compared to singlet T cells (T.90 signature) across T cell meta-clusters, and (B) 93 DEGs upregulated in in-contact monocytes compared to singlet monocytes (M.93 signature) across monocyte meta-clusters. For both A and B, paired per-participant points are shown for singlet (gold), coincidental (pink) and in-contact (blue) image-gated categories, with grey lines connecting matched participants; boxplots show the median and IQR. Image-gated categories were compared within each lineage by paired two-sided Wilcoxon signed-rank test with BH adjustment (adjusted p-values above brackets). Data were derived from 9,272 singlet, 12,022 coincidental and 12,487 in-contact T cells; 10,448 singlet, 11,002 coincidental and 11,915 in-contact monocytes from n=20 PBMC samples of patients with ATB collected at diagnosis.

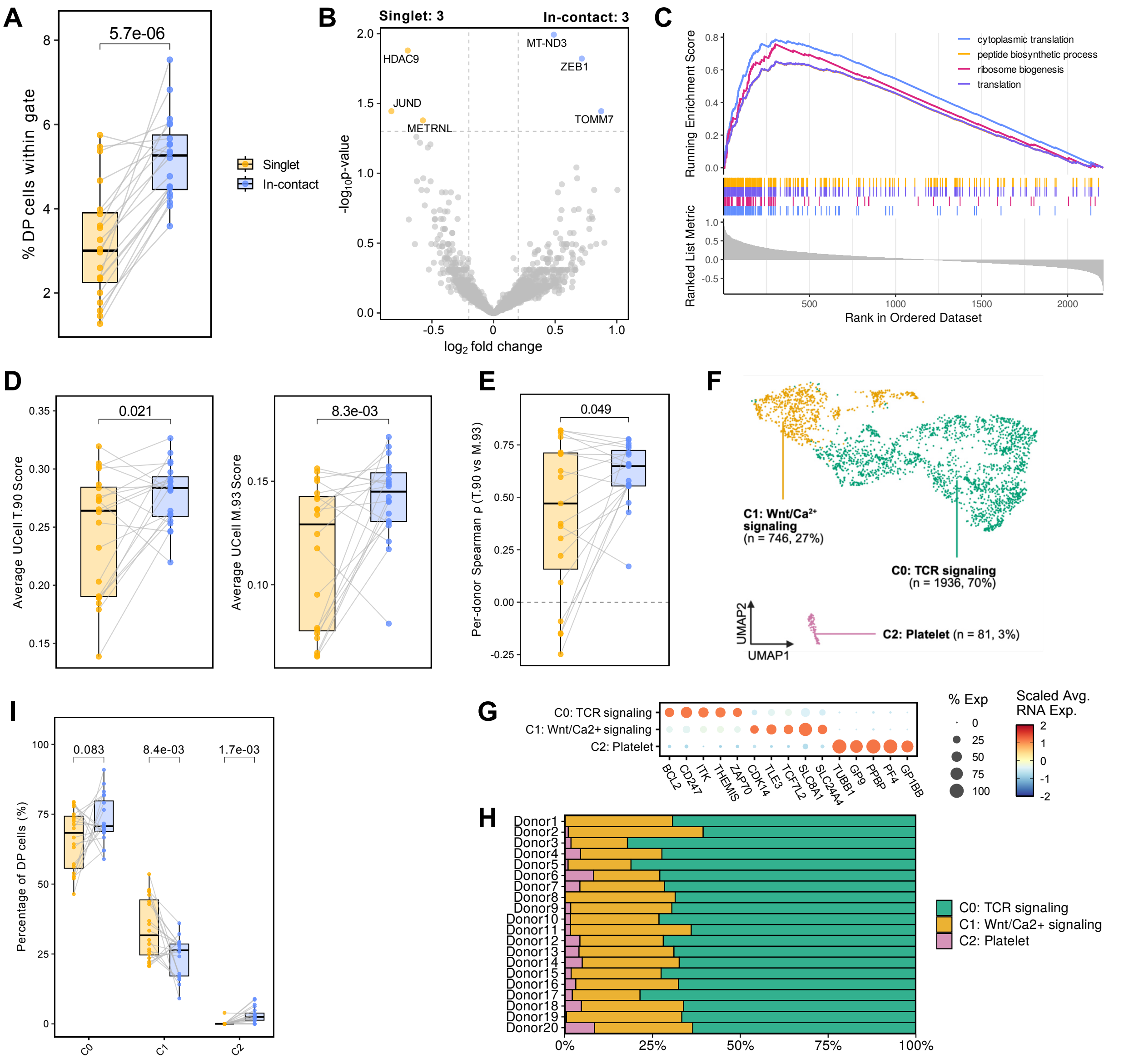


**Figure S7: Dual-lineage droplet analysis validates paired T cell-monocyte interaction signatures.** (A) Per-participant percentage of dual-lineage droplets within each gate (singlet, gold; in-contact, blue). The percentage was calculated per participant as the number of dual-lineage droplets divided by the total number of droplets in that gate. (B) Volcano plot of DEGs between in-contact (blue) and singlet (gold) dual-lineage droplets. Genes with BH-adjusted p < 0.05 and |average log2 FC| > 0.2 are colored by the gate in which they are upregulated. (C) GSEA enrichment score curves for the top four Gene Ontology:Biological Process pathways ranked by BH-adjusted p-value. The top panel displays the running enrichment score across the ranked gene list, the middle barcode panel marks the position of genes belong to the pathway, and the bottom panel indicates the ranking by log2FC from the DE analysis in Figure (B). (D) Boxplots of per-participant average UCell module scores for T.90 signature (left), and M.93 signature (right). (E) Boxplot of per-participant Spearman rank correlation coefficients ρ between the T.90 signature score and the M.93 signature score in singlet and in-contact categories. Overlapping genes between T.90 and M.93 signatures (n=63) were removed from the correlation analysis. (F) UMAP representation of dual-lineage droplets and manual cluster annotation. (G) Dot plot showing top markers genes for dual-lineage droplet clusters. Color intensity represents scaled average RNA expression levels; dot size corresponds to the percentage of cells within a given cluster expressing each gene. (H) Stacked bar plot showing the cell composition of dual-lineage droplet clusters per participant (normalized to 100%). (I) Dual-lineage droplet clusters composition differences. For A, D, E and I, paired per-participant points are shown for singlet (gold) and in-contact (blue) gates, with grey lines connecting matched participants; boxplots show the median and IQR. Gates were compared within each subset by paired two-sided Wilcoxon signed-rank test with BH adjustment (adjusted p-values above brackets). Data were derived from 597 singlet, 724 coincidental and 1,442 in-contact dual-lineage droplets from n=20 PBMC samples of patients with ATB collected at diagnosis.
